# Non-canonical role of a PHOSPHATE1 HOMOLOG 2 in suppressing seedling photomorphogenesis via the TOC1–PIF4 module

**DOI:** 10.64898/2026.05.09.724007

**Authors:** Devyan Das, Chirag Singhal, Bidhan Chandra Malakar, Sreeramaiah N. Gangappa

**Affiliations:** Department of Biological Sciences, Indian Institute of Science Education and Research Kolkata, Mohanpur 741246, West Bengal, India

**Keywords:** PHO1;H2, TOC1, PIF4, Gene regulation, Seedling Photomorphogenesis, Arabidopsis

## Abstract

Photomorphogenesis, the light-driven development of seedlings, is governed by a complex network of transcription factors and circadian regulators. While the TIMING OF CAB EXPRESSION 1 (TOC1) is known to link circadian rhythms with light-responsive growth, the mechanisms fine-tuning its activity remain poorly understood. Here, we identify PHOSPHATE 1 HOMOLOG 2 (PHO1;H2) as a novel negative regulator of seedling photomorphogenesis in *Arabidopsis*. Loss-of-function *pho1;h2* mutants exhibit hypersensitivity to light, characterized by markedly shorter hypocotyls and increased photopigment accumulation, whereas overexpression lines display reduced photomorphogenic response. We demonstrate that the N-terminal SPX domain of PHO1;H2 is both necessary and sufficient to repress seedling photomorphogenic growth. Mechanistically, *in vitro* and *in vivo* interaction assays reveal that the SPX domain physically binds and sequesters TOC1, inhibiting its regulatory function. This PHO1;H2-mediated sequestration of TOC1 alleviates the repression of PHYTOCHROME INTERACTING FACTOR 4 (PIF4), thereby promoting the expression of downstream genes involved in cell elongation and hormone signaling. Collectively, our findings reveal a competitive binding mechanism by which PHO1;H2 modulates the TOC1–PIF4 signaling axis, providing a crucial checkpoint for seedling growth in dynamic light environments.

## Introduction

Light is a key abiotic factor that determines plant growth and development (1–3). Plant development follows two morphogenic patterns, depending on the presence or absence of light: photomorphogenesis or skotomorphogenesis. Photomorphogenic growth is characterized by short hypocotyls, expanded cotyledons, and photopigment accumulation; in contrast, during skotomorphogenesis, seedlings exhibit long and pale hypocotyls with fused cotyledons, an apical hook and a lack of photopigments (4). Plants can sense even subtle changes in light quality, quantity, duration, and direction, leading to differential adaptations (5). For sensing these changes, plants possess different photoreceptors that detect specific wavelengths of light (6). These photoreceptors are classified into phytochromes, phototropins, and cryptochromes (7, 8). To perceive different wavelengths of light, plants have evolved to have five distinct classes of photoreceptors: red and far-red light are perceived by phytochromes (PHY A-E); blue light is perceived by cryptochromes (CRY1 and CRY2), phototropins (PHOT1 and PHOT2), as well as ZEITLUPE (ZTL)/ FLAVIN-BINDING, KELCH REPEAT, F-BOX 1 (FKF1)/ LOV kelch protein2 (LKP2); and UVR8 senses UV-B light (9).

Light signals activate these phytochromes, which, upon activation, translocate to the nucleus and interact with basic helix-loop-helix (bHLH) transcription factors, PHYTOCHROME INTERACTING FACTORS (PIFS) and inhibit them through phosphorylation and degradation (10, 11). PIFs control crucial developmental processes, including seed germination, de-etiolation, thermosensory growth, and flowering (12–14). The Phytochrome B (phyB), in the presence of red light, gets converted from its inactive (Pr) to active (Pfr) form and triggers the inhibition of PIFs and inactivation of CONSTITUTIVE PHOTOMORPHOGENESIS 1- SUPPRESSOR OF PHYA-105 (COP1-SPA) complex, leading to photomorphogenesis (15, 16). COP1, a RING E3 ubiquitin ligase, performs a central regulatory function, targeting several positive regulators of photomorphogenesis for degradation in darkness, including ELONGATED HYPOCOTYL 5 (HY5) (17–22). DE-ETIOLATED 1 and SUPPRESSOR OF PHYA-105 (SPA1) are other E3 ubiquitin ligases that function in association with COP1 and promote HY5 degradation (23, 24).

The PHYTOCHROME INTERACTING FACTOR 4 (PIF4) interacts with circadian clock elements to maintain optimum photomorphogenesis and promote thermomorphogenesis (25–28). In Arabidopsis, CCA1, LHY, and TOC1 are considered the central oscillators of the circadian clock, and recent studies have shown that TOC1 interacts with and inhibits PIF4 to mediate the circadian gating of thermoresponsive growth (28). During the day, when there is a surge in temperature, *CIRCADIAN CLOCK ASSOCIATED1* (*CCA1*) (a morning-phased transcription factor)/ LATE ELONGATED HYPOCOTYL (CCA1/LHY) positively regulate the *PIF4* expression (29, 30). TIMING OF CAB EXPRESSION1 (TOC1) accumulates in the evening and physically interacts with PIF4, inhibiting its function and gating thermoresponsive growth to earlier in the day (28). Parallelly, PSEUDORESPONSE REGULATOR 5 (PRR5), PRR7, and TOC1 inhibit PIF4 function, both at the transcriptional level and through sequestration, ensuring tight and temporal control of PIF4 activity (31). The Evening Complex (EC), consisting of EARLY FLOWERING3 (ELF3), ELF4 and LUX ARRHYTHMO (LUX) (ELF3/ELF4/LUX), binds to the *PIF4* promoter, repressing its transcription and inhibiting hypocotyl elongation (26). ELF3 further suppresses PIF4 activity through direct physical interaction, independent of the EC, thereby inhibiting its function (25).

The Arabidopsis PHOSPHATE 1 HOMOLOG (PHO1) protein family, characterized by Conserved SPX (SYG1/Pho81/XPR1) and EXS (ERD1/XPR1/SYG1) domains, comprises 11 members with a conserved structural topology featuring an N-terminal SPX and C-terminal EXS domain (32, 33). The SPX domain functions as a sensor for inositol pyrophosphates (InsPs), regulating phosphate homeostasis and contributing to light signaling. In contrast, the EXS domain is implicated in ER protein localization (34). SHORT HYPOCOTYL UNDER BLUE 1 (SHB1), a PHO1 family member (PHO1;H4), promotes PIF4-mediated growth by acting as a transcriptional co-activator of CCA1, enhancing *PIF4* expression around midday to facilitate thermosensory hypocotyl elongation in response to warm ambient temperature and light intensity (29, 35). While the role of SHB1 (PHO1;H4) in light- and temperature-mediated growth regulation is well-established, the functions of other PHO1 homologs remain unexplored. Using a reverse genetic approach, we identified PHO1;H2 as a negative regulator of seedling photomorphogenesis in Arabidopsis. Mutants lacking *PHO1;H2* (*pho1;h2*) exhibit a hyperphotomorphogenic response with enhanced hypocotyl growth inhibition and increased photopigment accumulation under white light and various wavelengths of light. Conversely, *PHO1;H2* overexpression lines displayed elongated hypocotyls and reduced photopigment levels. *PHO1;H2* expression is reduced in light, while it peaks in the dark. We further demonstrated that PHO1;H2 interacts with TIMING OF CAB EXPRESSION 1 (TOC1) via its SPX domain, both in vitro and in vivo, alleviating TOC1-mediated repression of PIF4 transcriptional activity. Consistent with this, *PHO1;H2* overexpression enhanced, while *pho1;h2* mutation suppressed, the long-hypocotyl phenotype in the *PIF4* overexpression line. Together, this study uncovers a novel role and the molecular mechanism by which PHO1;H2 suppresses seedling photomorphogenesis, highlighting its distinct function within the PHO1 family.

## Results

### PHO1;H2 is a homolog of the PHO1 family of proteins, and is highly conserved across plant species

The PHO1 family of proteins comprises multiple homologs (PHO1;H1-PHO1;H10) and has been widely studied for inorganic phosphate transport and homeostasis in plants (36, 37). Among them, PHOSPHATE1 HOMOLOG 4, also known as SHORT HYPOCOTYL UNDER BLUE1 (SHB1), has previously been implicated in regulating seed development (33), seedling growth, and thermosensory responses via its interactions with CCA1 (29, 35), indicating that the PHO1 homologs have a broader role in seedling morphogenesis beyond phosphate transport. In this study, we identified and functionally characterized another homologue of PHO1, i.e. PHO1;H2. Phylogenetic analysis and domain architecture reveal that the PHO1 homolog group of proteins shares an evolutionarily conserved sequence among plant genomes (32, 38). The protein sequence of AtPHO1;H2 was aligned and compared with diverse plant genomes. We found that PHO1;H2 is highly conserved across the seedless and dicot species included in the study (Fig. S1A). Interestingly, AtPHO1;H2 shows an E-value of 0 in dicots, indicating a nearly identical sequence. Monocots, with low E-value, still show significant conservation, suggesting greater sequence divergence between monocots and dicots. (Fig. S1A).

To further substantiate these results, we aligned and compared the amino acid sequences of AtPHO1;H2 with those of representative land plant species, including monocots and seedless plants, using NCBI’s COBALT multiple sequence alignment tool, revealing a topology consistent with established plant evolutionary relationships (Fig. S1B). The tree clearly resolves three clades: seedless, monocots, and dicots. Seedless plants, *Selaginella moellendorffii* (a lycophyte) and *Sphagnum fallax* (a moss), branched earlier than both monocots and dicots, indicating an ancient origin of PHO1;H2, predating the divergence of seed plants (Fig. S1B). The longer branch length of seedless plants suggests greater evolutionary divergence from angiosperms. Further, the tree clearly resolves the two angiosperm clades, dicots and monocots. Dicots form a monophyletic clade, and the branch lengths within the dicots were relatively short, showing tight clustering of related taxa (e.g., *Arabidopsis* and *Brassica*; *Solanum* species together), indicating minimal divergence (Fig. S1B). However, slightly longer lengths in grasses suggest that the protein has diverged moderately in monocots than in dicots. We also found that the N-terminal SPX and C-terminal EXS domains are highly conserved across all the examined plant genomes (Fig. S1), suggesting a fundamental and likely conserved role of PHO1;H2 across species.

### PHO1;H2 is a negative regulator of photomorphogenesis

To understand the role of PHO1;H2 in seedling photomorphogenesis, we obtained two independent T-DNA insertion mutant lines from the Eurasian Arabidopsis Stock Centre (uNASC). The *PHO1;H2* gene consists of 12 exons. In the *pho1;h2-1* mutant, the T-DNA insertion is in the third exon; in the *pho1;h2-2* mutant, it is in the 10^th^ exon (Fig. S2A-C). Analysis for the transcript accumulation in these mutant alleles revealed no transcript accumulation in the *pho1;h2-1* and *pho1;h2-2* alleles, suggesting that both alleles are likely null (Fig. SA-C). To examine the role of PHO1;H2 in *Arabidopsis* seedling photomorphogenesis, we grew two independent alleles, *pho1;h2-1* and *pho1;h2-2,* along with the Col-0, in white-light (WL) under short-day (SD, 8h light/16h) photoperiod and measured the hypocotyl length of six-day-old seedlings. Our data suggest that both *pho1;h2-1* and *pho1;h2-2* alleles displayed significantly shorter hypocotyls than Col-0 under WL across different fluences tested (Fig. 1A-C), suggesting that PHO1;H2 functions as a negative regulator of light-mediated inhibition of seedling hypocotyl growth. Next, to check if PHO1;H2-mediated suppression of inhibition of seedling hypocotyl growth is specific to any particular wavelength of light, we grew Col-0 and *pho1;h2* mutants in red-light (BL), blue-light (RL), and far-red light (FRL) for six days and measured their hypocotyl lengths. Our data suggest that *pho1;h2* mutant alleles showed significantly shorter hypocotyls than the Col-0 across all the fluence rates of BL (D-F), RL (G-I), and FRL (Fig.1J-L), indicating that PHO1;H2 suppresses the inhibition of seedling hypocotyl growth in a wavelength-independent manner.

**Fig. 1.**
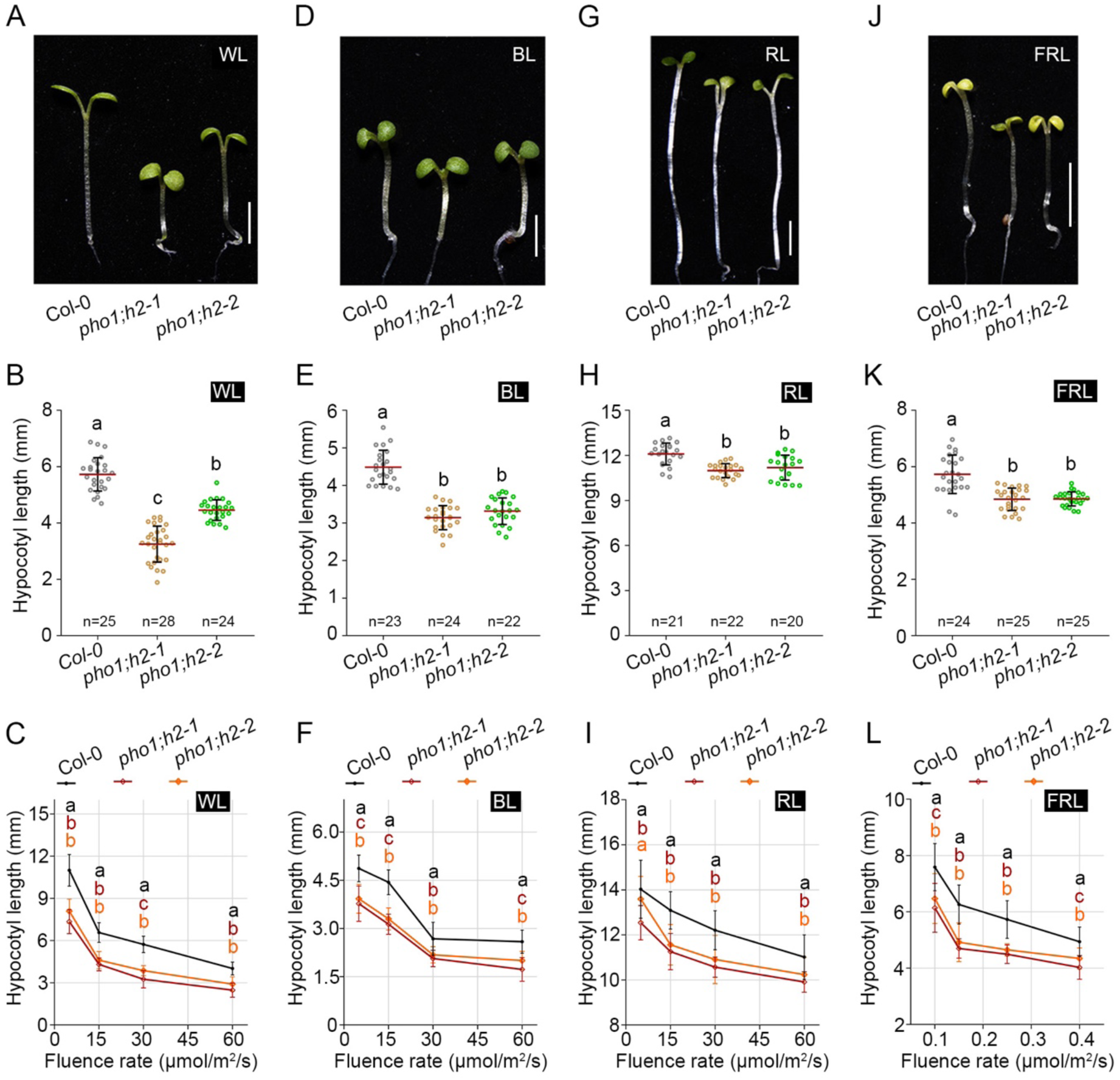
PHO1;H2 is a negative regulator of seedling photomorphogenesis in Arabidopsis. (**A-D**) Representative hypocotyl images of six-day-old seedlings of Col-0 and *pho1;h2* mutants (*pho1;h2-1* and *pho1;h2-2*) grown under short days at 22°C in various lights: White Light (30 µmol m²s¹) (A), Red Light (30 µmol m²s¹) (B), Blue Light (30 µmol m²s¹)(**C**) and Far-red Light (0.41 µmol m²s¹) (**D**). Scale = 1mm. (**E-H**) Mean hypocotyl length of Col-0 and *pho1;h2-1, pho1;h2-2* mutants under different wavelengths of light WL (A), BL (B), RL (C) and FRL (D). The scatter dot plot shows the data points for each genotype; the error bars show the standard deviation, and the red line marks the mean. The asterisk (*) represents the statistically significant differences from Col-0 and between the indicated pair of genotypes as revealed by Student’s *t*-test (\**p*<0.05; \*\**p*<0.001; ***p<0.001; ****p<0.0001). “n” denotes the number of seedlings measured. (**I-L**) Hypocotyl growth rate curve of Col-0 and *pho1;h2* mutants under increasing fluence of WL (5,15, 30, 60) µmol m-²s-¹ (**I**), BL (5, 15, 30, 60) µmol m-²s-¹ (**J**), RL (5, 15, 30, 60) µmol m-²s-¹ (K) and FRL (0.1,0.15,0.2,0.4) µmol m-²s-¹ (L). Small letters above the bars denote genotypes that significantly differ as revealed by one-way ANOVA followed by Tukey’s post-hoc HSD test (p<0.05). “*n*” indicates the number of seedlings measured. (**M** and **N**) RT-qPCR data showing transcript abundance of growth-promoting genes such as *IAA19*, *SAUR22,* and *YUC8,* and light-inducible genes *RBCS1A* and *CAB1.* The six-day-old seedlings of Col-0 and *pho1;h2-1* were harvested under SD conditions for gene expression analysis, with *TUB2* as an endogenous control. Relative gene expression was calculated by normalizing to Col-0 (as 1). The error bar shows the standard deviation (n =3 biological replicates). Small letters above the bars denote genotypes that significantly differ as revealed by one-way ANOVA followed by Tukey’s post-hoc HSD test (p<0.05).

### PHO1;H2 inhibits the expression of de-etiolation marker genes while promoting the expression of growth genes

Hypocotyl elongation is facilitated by cell elongation, which is achieved by loosening the cell wall via modulation of auxin biosynthesis and signaling (27, 39–41). Because the *pho1;h2* mutant exhibits reduced hypocotyl length compared to Col-0, we examined the expression of growth-related genes. Our RT-qPCR analysis revealed that growth-responsive genes such as *XTR7 (XYLOGLUCAN ENDOTRANSGLUCOSYLASE/HYDROLASE 7), EXP8 (EXPANSIN A8), IAA19 (INDOLE-3-ACETIC ACID INDUCIBLE 19*)*, SAUR22* (*SMALL AUXIN UP RNA 22*) and *YUC8* (*YUCCA8)* showed robust downregulation in the *pho1;h2-1* mutant compared to Col-0 (Fig. 2A-E). We also examined the expression of light-responsive genes involved in the photosynthetic machinery and photopigment accumulation, including those involved in chlorophyll and anthocyanin biosynthesis (42, 43). Interestingly, unlike growth-promoting genes, light-inducible genes such as *RBCS1A* (*RIBULOSE BISPHOSPHATE CARBOXYLASE SMALL CHAIN 1A*)*, CAB1* (*CHLOROPHYLL A/B BINDING PROTEIN 1*) and *ELIP2* (*EARLY LIGHT INDUCED PROTEIN 2*) that are part of photosynthetic machinery showed significant upregulation in the *pho1;h2-1* mutant compared to Col-0 (Fig. 2F-H). Similarly, the anthocyanin biosynthetic genes*, CHALCONE ISOMERASE* (*CHI*) and *CHALCONE SYNTHASE* (*CHS*), also showed upregulation in the *pho1;h2-1* mutant compared to Col-0 (Fig. 2I,J). These results indicate that the *pho1;h2* mutant phenotype is likely due to reduced expression of growth-promoting genes and enhanced expression of de-etiolation marker genes.

**Fig. 2.**
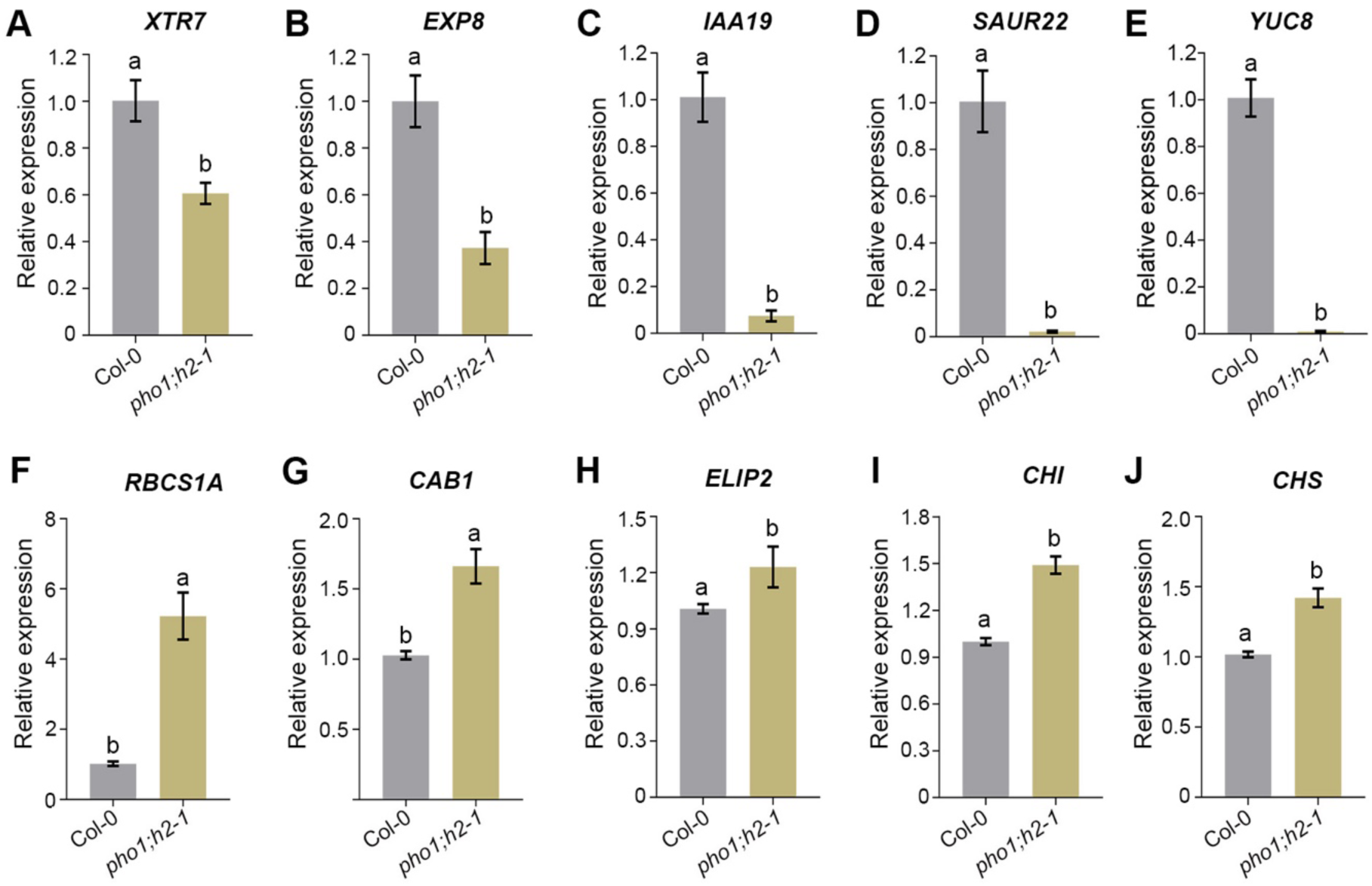
PHO1;H2 inhibits the expression of de-etiolation marker genes while promoting the expression of growth genes. **(A-E)** RT-PCR analysis of growth-related genes *XTR7, EXP8, IAA19*, *SAUR22,* and *YUC8* in Col-0 and *pho1;h2-1* mutant. **(F-J)** Gene expression analysis of light-inducible genes *RBCS1A*, *CAB1, ELIP2* (H), *CHI* (I) and *CHS* (J) in col-0 and *pho1;h2-1.* The six-day-old seedlings of Col-0 and *pho1;h2-1* were harvested at ZT4 under SD conditions for gene expression analysis, with *TUB2* as an endogenous control. Relative expression was calculated with respect to the gene expression of the *pho1;h2-1* mutant. The error bar shows the SD (n=3 biological replicates). Small letters above the bars denote genotypes that significantly differ as revealed by one-way ANOVA followed by Tukey’s post-hoc HSD test (p<0.05).

### The *PHO1;H2* gene expression is controlled by light

As PHO1;H2 regulates light-mediated seedling development, we wanted to examine if light controls its spatial-temporal gene expression. We grew Col-0 seedlings under SD (8 h light/16 h dark, 60 μmol m^-2^ s^-1^) and harvested the tissue at different ZT time points throughout the day, followed by mRNA extraction and RT-qPCR analysis. Our data reveal that the *PHO1;H2* transcript accumulation exhibits a diurnal expression pattern, wherein the *PHO1;H2* transcript gradually decreases from ZT0 to ZT6, while it starts increasing its accumulation gradually through the night from ZT10 to ZT24, and reaches a maximum at ZT24 (Fig. S3A), indicating that the expression of *PHO1;H2* is repressed by light and induced by dark. To further validate this, we conducted both light-to-dark and dark-to-light shift experiments. When six-day-old continuous dark-grown seedlings were transferred to the light for indicated time periods (hours), such as 0, 2, 4, 8, 16 and 24h, we found that the *PHO1;H2* accumulation is significantly decreased at 2h light treatment than the 0h control, and it showed further decline from ZT2-ZT24 (Fig. S3B). However, when six-day-old continuous light seedlings were shifted to darkness for different time intervals as indicated, we found that *PHO1;H2* expression was significantly increased at a 2h dark treatment compared to a 0h control (Fig. S3C). Furthermore, its expression showed a gradual and significant increase, reaching a maximum at 24h of dark treatment (Fig. S3C). Together, these RT-qPCR results confirm that light suppresses *PHO1;H2* transcription, while dark treatment promotes it.

We also investigated the transcript abundance of *PHO1;H2* across various aerial tissues of mature Col-0 plants. Col-0 plants were grown under short-day conditions, and after 45 days post-germination, the tissue was harvested from rosette leaves, cauline leaves, siliques and flowers. RNA was extracted from each tissue, and qPCR was conducted with *EF1α* as an internal control. Our data revealed that the *PHO1;H2* transcript is markedly higher in cauline leaves and flowers compared to rosette leaves and siliques (Fig. S3D).

### PHO1;H2 physically interacts with TOC1 through its N-terminal SPX domain

Mutants of *PHO1;H2* displayed a short hypocotyl phenotype, and gene expression analysis revealed a rhythmic transcript abundance. Previous findings reported that the SHB1 physically interacts with CCA1 through its N-terminal SPX domain to regulate hypocotyl growth in response to light and temperature cues. (29, 35). To reveal potential molecular interactions of PHO1;H2 with any of the circadian clock proteins, we performed protein-protein interaction studies using the Yeast two-hybrid (Y2H) assay. We cloned core circadian clock genes such as *ELF3* (*EARLY FLOWERING 3*)*, ELF4* (*EARLY FLOWERING 4*)*, PRR5 (PSEUDO-RESPONSE REGULATOR 5), PRR7 (PSEUDO-RESPONSE REGULATOR), CCA1 (CIRCADIAN CLOCK ASSOSCIATED 1),* and *TOC1 (TIMING OF CAB EXPRESSION1)* in pGADT7 vector containing GAL4 Activating domain (AD), and PHO1;H2 was cloned in pGBKT7 vector containing GAL4-DNA binding domain (DBD). The Y2H results revealed that PHO1;H2 could physically interact with TOC1, but not with the other clock proteins examined in our Y2H assays (Fig. S4). Interestingly, TOC1, a central component of the evening-phase clock regulator, inhibits PIF4 function by sequestering it, thereby regulating hypocotyl growth (28). PHO1;H2 is composed of two distinct functional domains, the N-terminal SPX and the C-terminal EXS domains. To know which domain of PHO1;H2 is responsible for the interaction with TOC1, we amplified the N-terminal SPX domain (1-348 amino acids) and the C-terminal EXS domain (607-807 amino acids), separately. We cloned them into pGBKT7, which contains a GAL4-binding domain, to serve as bait. We co-transformed TOC1-AD (prey) with PHO1;H2-SPX-BD (SPX-BD) or PHO1;H2-EXS-BD (EXS-BD) onto Y2H-Gold reporter strain, and screened on synthetic media lacking Leucine, Tryptophan, Histidine and Adenine (quadruple drop-out, QDO) and in the presence of 3-amino-1,2,4-triazole (3-AT) (Fig. 3A). The co-transformants of TOC1-AD with SPX-BD were able to grow on the minimal QDO media. However, the TOC1-AD with EXS-BD co-transformants failed to grow on minimal QDO media, as did the negative controls (Fig. 3A), suggesting that the PHO1;H2 interaction with TOC1 is mediated by the SPX domain rather than the EXS domain. The T antigen and p53 combination served as a positive control (Fig. 3A). Together, these results indicate that the SPX domain is both necessary and sufficient for the interaction with TOC1 (Fig. 3A).

**Fig 3.**
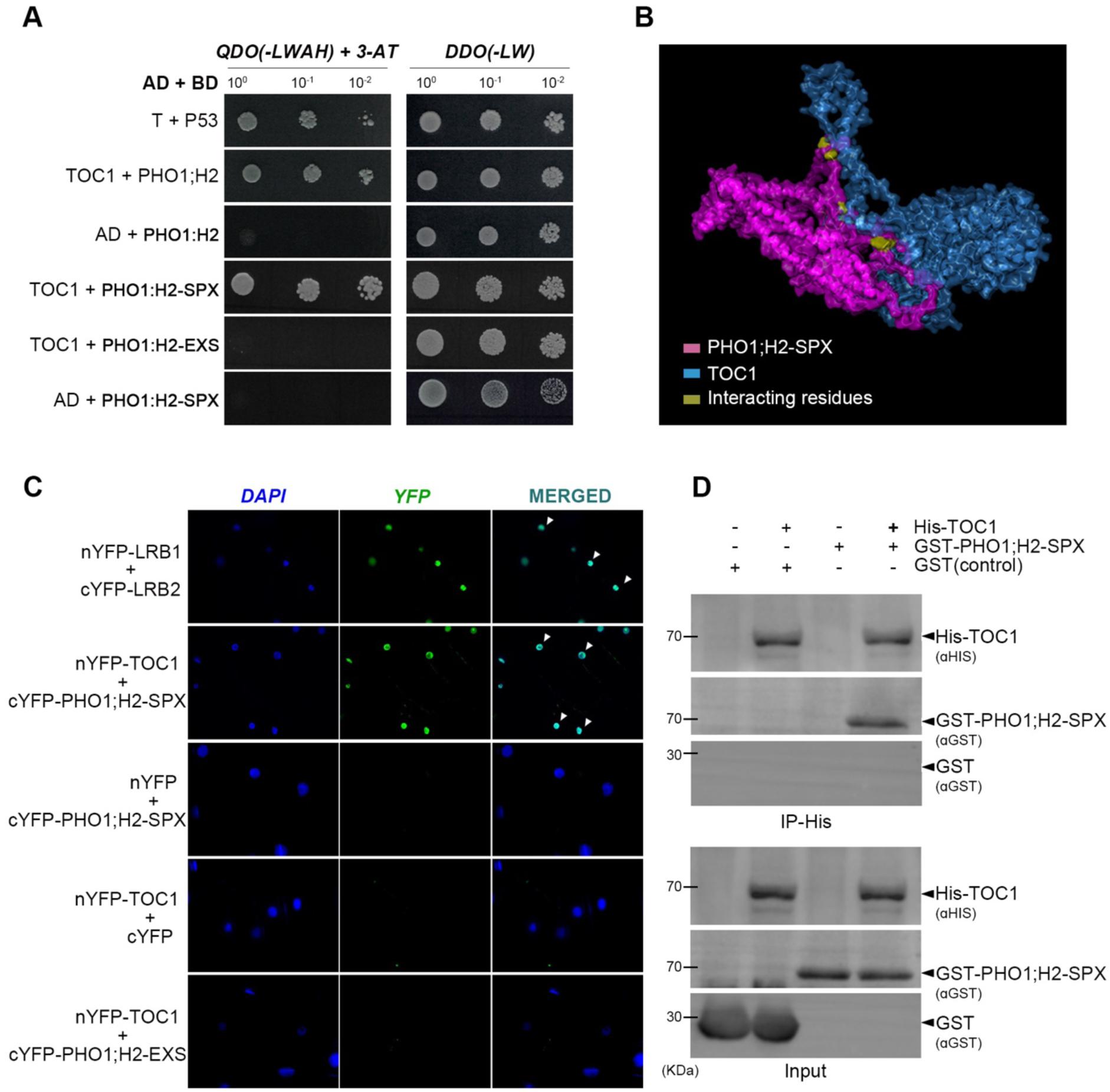
PHO1;H2 physically interacts with TOC1 through its N-terminal SPX domain. (**A**) *in-silico* molecular docking of modelled structure of PHO1;H2-SPX domain and TOC1 using *I-Tasser*. The docked structure is represented as a surface. Common residues from full-length PHO1;H2 and PHO1;H2-SPX interacting with TOC1 have been highlighted. (**B**) Yeast two-hybrid assays showing the physical interaction between PHO1:H2 and PHO1;H2-SPX with TOC1. Co-transformants were first selected on double dropout medium (DDO, SD/-Leu/-Trp) to confirm the presence of both bait and prey constructs (left panel), and subsequently cultures were dropped onto quadruple dropout medium (QDO, SD/-Ade/-His/-Leu/-Trp) together with 3-AT to assess protein–protein interactions (right panel). (**C**) *In-vitro* pull-down assay performed with TOC1-His as bait (pulled down using Ni-NTA beads), and in the pulled-down sample, GST-SPX was detected using anti-GST antibody through immunoblotting. (**D**) BIFC assay, shows the physical interaction between nYFP-TOC1+cYFP-PHO1;H2 and nYFP-TOC1+cYFP-PHO1;H2-SPX in onion epidermal cells. The nYFP-LRB1 and cYFP-LRB2 combination was used as a positive control. Scale = 5 µm.

To further explore, which residues are responsible for the interaction between PHO1;H2, and TOC1, we employed a structure-based protein-protein interaction approach. PHO1;H2, and TOC1 were modelled using the I-TASSER server, generating reliable 3D structures for docking. Prior evidence from the Y2H assay revealed that the SPX domain of PHO1;H2 interacts with TOC1. We modelled the SPX domain separately and docked SPX and full-length PHO1;H2 against TOC1 using CLUS-PRO (Fig. 3B). The resultant docked complexes (PHO1;H2-FL-TOC1 and PHO1;H2-SPX-TOC1) were analyzed using the PDBsum server to identify interfacial residues involved in the interaction (Fig. S5A,B). Notably, the comparative analysis revealed a set of common interfacial residues present in both complexes: Ala210, Ala212, Arg209, Arg224, Asn207, Glu225, His213, Phe222, Thr205, and Tyr61 (Fig. S5C). All these residues localize to the N-terminal SPX domain of PHO1;H2, reinforcing the above results. Further quantitative analysis of the complexes using the PRODIGY web server revealed a significant difference in binding thermodynamics (Fig. S5D). The PHO1;H2-TOC1 exhibited a binding free energy (ΔG) of -25 kcal/mol, whereas SPX-TOC1 displayed a similar but slightly more favorable (ΔG) of -25.6 kcal/mol, and the dissociation constant for the SPX-TOC1 complex was lower (1.8×10^-19^ M) than PHO1;H2-TOC1 (4.5 ×10^-19^ M) (Fig. S5D), suggesting that SPX has a higher binding affinity to TOC1. Together, the yeast two-hybrid and molecular docking analysis confirm that the SPX domain of PHO1;H2 is necessary and sufficient to mediate the interaction with TOC1.

These protein-protein interactions were further confirmed both in vitro pull-down assays and in planta. First, we carried out in planta protein-protein interaction in onion epidermal cells using the Bimolecular Fluorescence Complementation (BiFC) assay. We cloned *PHO1;H2* onto the *pSPYCE* vector with a C-terminal YFP tag (*PHO1;H2-cYFP*), and *TOC1* onto the *pSPYNE* vector with an N-terminal YFP tag (*TOC1-nYFP*). When *PHO1;H2-cYFP* and *TOC1-nYFP* constructs were co-infiltrated into the onion epidermal cells through *Agrobacterium tumefaciens* (GV3101) (44), a strong YFP signal was detected in the nucleus after three days of incubation in the dark. However, no signal was detected in the samples infiltrated with *Agrobacterium* containing *PHO1;H2-cYFP* and empty *pSPYNE* (nYFP) vector, indicating that PHO1;H2-cYFP and TOC1-nYFP likely interact in planta (Fig. 3C). Interestingly, as seen above, when the *PHO1;H2-SPX-cYFP* construct was co-infiltrated together with *TOC1-nYFP*, it also showed a strong YFP signal in the nucleus (Fig. 3C), while the *PHO1;H2-EXS-cYFP* construct together with *TOC1-nYFP* failed to reconstitute YFP fluorescence (Fig. 3C). Next, we performed the in vitro pulldown assay using *Escherichia coli (E. coli)* purified recombinant His-tagged TOC1 (TOC1-His, used as bait) and GST-tagged PHO1;H2-SPX (GST-PHO1;H2-SPX, used as prey). When His-TOC1 was pulled down using Ni-NTA beads, we could detect GST-PHO1;H2-SPX protein in the co-immunoprecipitated sample, using anti-GST antibody (Fig. 3D). Immunoblot analysis revealed the presence of PHO1;H2-SPX in the eluate from the reaction containing His-TOC1 and PHO1;H2-SPX, while we did not detect GST in a reaction containing His-TOC1 along with only GST (used negative control) (Fig. 3D). Together, data from multiple lines of evidence confirm that PHO1;H2 physically interacts with TOC1 via its N-terminal SPX domain (Fig. 3D).

### PHO1;H2 genetically interacts with TOC1 and likely inhibits its function

Based on the above findings, we were curious to understand the genetic interactions between PHO1;H2 and TOC1. We generated the homozygous double mutant *pho1;h2-1 toc1-2* and measured hypocotyl length of parental and Col-0 genotypes grown in WL under SD for six days. Our data reveal that the *toc1-2* mutant exhibited significantly longer hypocotyls than the wild type (Fig. 4A,B), as reported (45), underscoring that TOC1 promotes photomorphogenesis. Interestingly, the long hypocotyl phenotype of the *toc1-2* was significantly suppressed by *pho1;h2-1*, suggesting that the *toc1-2* hypocotyl phenotype is to a greater extent dependent on PHO1;H2 (Fig. 4A,B). As PIF4 is the key promoter of hypocotyl elongation and is also repressed by TOC1 (28), we wanted to determine whether this repression involves altered PIF4 protein levels. We performed immunoblotting to assess PIF4 abundance in six-day-old SD-grown seedlings of Col-0, single- and double-mutant genotypes, along with *pifQ* (used as a negative control). Immunoblot analysis using anti-PIF4 antibodies reveals that PIF4 abundance in *pho1;h2-1*, *toc1-2* and *pho1;h2-1 toc1-2* double mutant genotypes is comparable to Col-0, suggesting the defect in hypocotyl growth is not attributable to PIF4 protein abundance (Fig. 4C)

**Fig 4.**
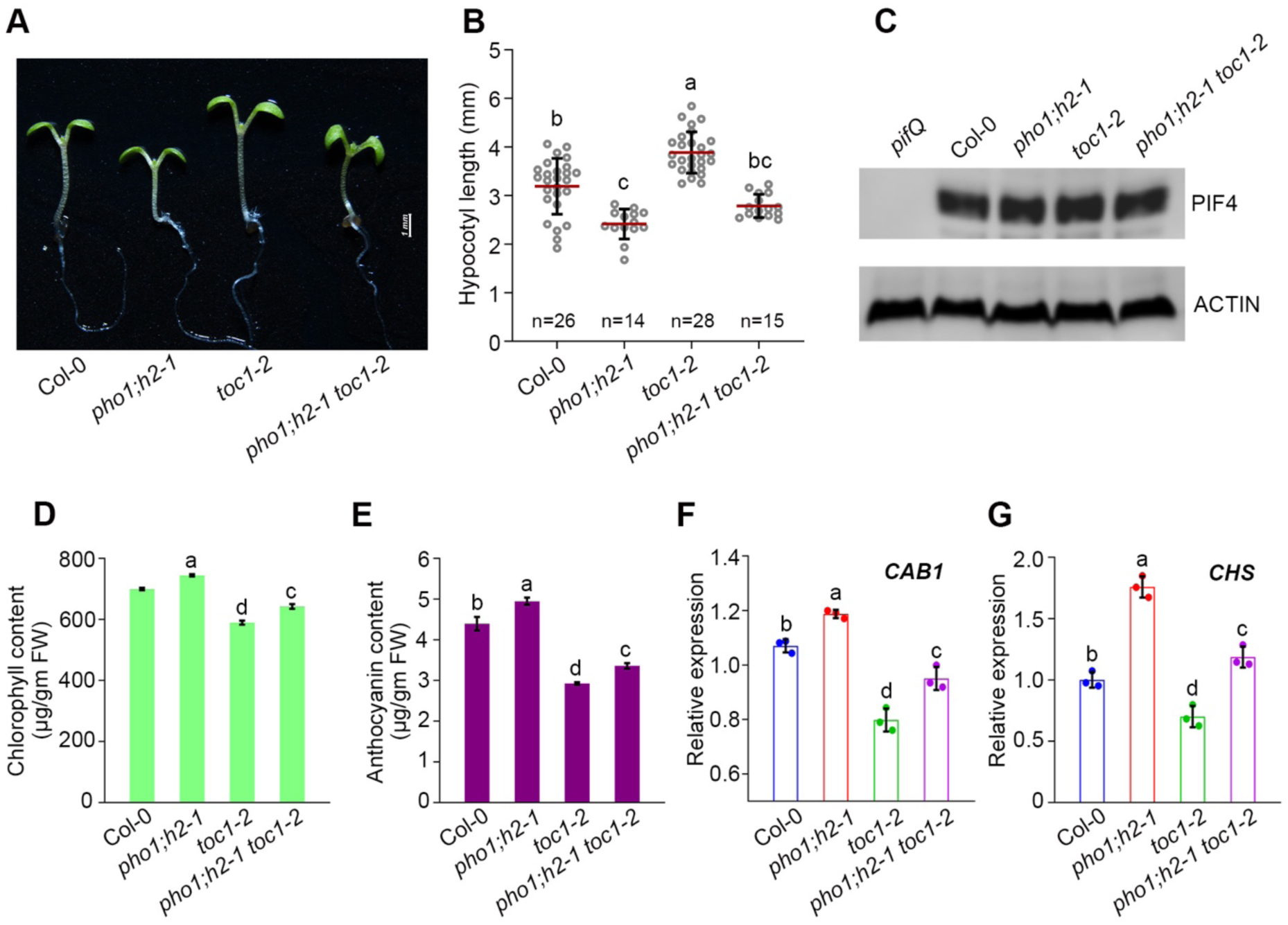
PHO1;H2 genetically interacts with TOC1 and inhibits its function. **(A)** Representative hypocotyl images of six-day-old seedlings of Col-0*, pho1;h2-1*, *toc1-2* and *pho1;h2-1 toc1-2* grown in WL (60 µmol m-²s-¹) under SD. Scale = 1 mm. (B) Hypocotyl length measurements for the seedlings shown in (A). The scatter dot plot shows the data points for each genotype; the error bars show the standard deviation, and the red line marks the mean. Small letters above the bars denote genotypes that significantly differ as revealed by one-way ANOVA followed by Tukey’s post-hoc HSD test (p<0.05). “*n”* indicates the number of seedlings measured. (C) Immunoblotting with native anti-PIF4 antibodies showing PIF4 protein levels in Col-0*, pho1;h2-1, toc1-2* and *pho1; h2-1 toc1-2* double mutant are unaltered. Six-day-old seedlings grown under SD in WL were used for the immunoblotting. The *pifQ* mutant was used as a negative control. Actin levels indicate equal protein loading. (**D** and **E**) Quantification of chlorophyll (D) and anthocyanin (E) content in six-day-old seedlings of Col-0*, pho1;h2-1, toc1-2* and *pho1;h2-1 toc1-2* grown in WL (60 µmol m^-^²s^-^¹) under SD photoperiod. Error bar denotes the standard deviation for n=3. Small letters above the bars denote genotypes that significantly differ as revealed by one-way ANOVA followed by Tukey’s post-hoc HSD test (p<0.05). *n* indicates the number of seedlings measured. (**F** and **G**) RT-qPCR data showing transcripts of light-inducing genes *CAB1* (F) and *CHS* (G) from six-day-old seedlings of Col-0, *pho;h2-1, toc1-2* and *pho1;h2-1 toc1-2* genotypes grown in 22°C WL under SD conditions. Tissue was harvested at ZT4 under SD. *TUB2* was taken as an endogenous control. The asterisk (*) represents the statistically significant differences from Col-0 and between the indicated pair of genotypes as revealed by Student’s *t*-test (\**p*<0.05; \*\**p*<0.001; ***p<0.001; ****p<0.0001).

Photomorphogenic development of seedlings is characterized by increased chlorophyll and anthocyanin accumulation (46). We quantified chlorophyll and anthocyanin content in the six-day-old *pho1*;*h2 toc1-2* mutant seedlings and compared it with Col-0. The chlorophyll and anthocyanin content were reduced in the *toc1-2* mutant, while the photopigment content was elevated in the *pho1;h2-1* mutant compared to the wild type (Fig. 4D,E). In the *pho1*;*h2 toc1-2* double mutant background, the photopigment level was strongly suppressed compared to the *pho1;h2-1* (Fig. 4D,E). The photopigment levels were significantly lower than in Col-0 and closer to those of the *toc1-2* mutant (Fig. 4D,E). Concomitantly, the expression of key light-responsive genes, such as *CAB1* and *CHS,* was significantly elevated in the *pho1*;*h2-1* mutant, while downregulated in the *toc1-2* mutant, compared to Col-0 (Fig. 4F,G). However, *CAB1* and *CHS* expression in the *pho1*;*h2-1 toc1-2* double mutant was significantly suppressed compared to the *pho1*;*h2* single mutant (Fig. 4F,G), suggesting that both PHO1;H2 and TOC1 exert antagonistic effects in controlling the expression of light-responsive genes and seedling photomorphogenic responses. The intermediate level of photopigment and gene expression in *pho1;h2-1 toc1-2* double mutant supports that PHO1;H2 and TOC1 pathways converge to modulate hypocotyl elongation without affecting PIF4 protein accumulation.

### SPX domain of PHO1;H2 is necessary and sufficient to suppress photomorphogenesis

The above results suggest that PHO1;H2 physically and genetically interacts with TOC1 and functions antagonistically to regulate seedling photomorphogenic growth. As the N-terminal SPX domain of PHO1;H2 mediated its physical interaction with TOC1, we wanted to test whether the SPX domain alone is capable of suppressing photomorphogenesis. We generated transgenic lines overexpressing, the full-length (FL) PHO1;H2 (807 amino acids), the N-terminal SPX (1-348 amino acids) and the C-terminal EXS (607-807 amino acids) domains, separately (Fig. 5A). The homozygous transgenics lines with increased *PHO1;H2* expression (Fig. S6A-C) were used to measure the hypocotyl length from six-day-old seedlings grown under SD. Transgenic lines overexpressing FL PHO1;H2 (*35:PHO1;H2*-Myc #1 and *35S:PHO1;H2-Myc #2*) showed significant elongation in hypocotyl growth compared to Col-0 (Fig. 5B,C). Interestingly, transgenics lines overexpressing the *PHO1;H2-SPX* (*35S:PHO1;H2-SPX-Myc #1* and *35S:PHO1;H2-SPX-Myc #2*) exhibited significantly longer hypocotyls compared to Col-0 seedlings (Fig. 5B,C). However, the transgenic lines overexpressing only the EXS domain, C-terminal part of *PHO1;H2* (*35S:PHO1;H2-EXS-Myc #1* and *35S:PHO1;H2^-^EXS-Myc #2*) did not show any significant difference in the hypocotyl length compared to the Col-0 seedlings (Fig. 5B,C). Notably, the hypocotyl length between *35S:PHO1;H2-Myc* and *35S:PHO1;H2-SPX-Myc* transgenic lines was comparable, indicating that the SPX domain of PHO1;H2 is necessary and probably sufficient for suppressing seedling photomorphogenesis.

**Fig 5.**
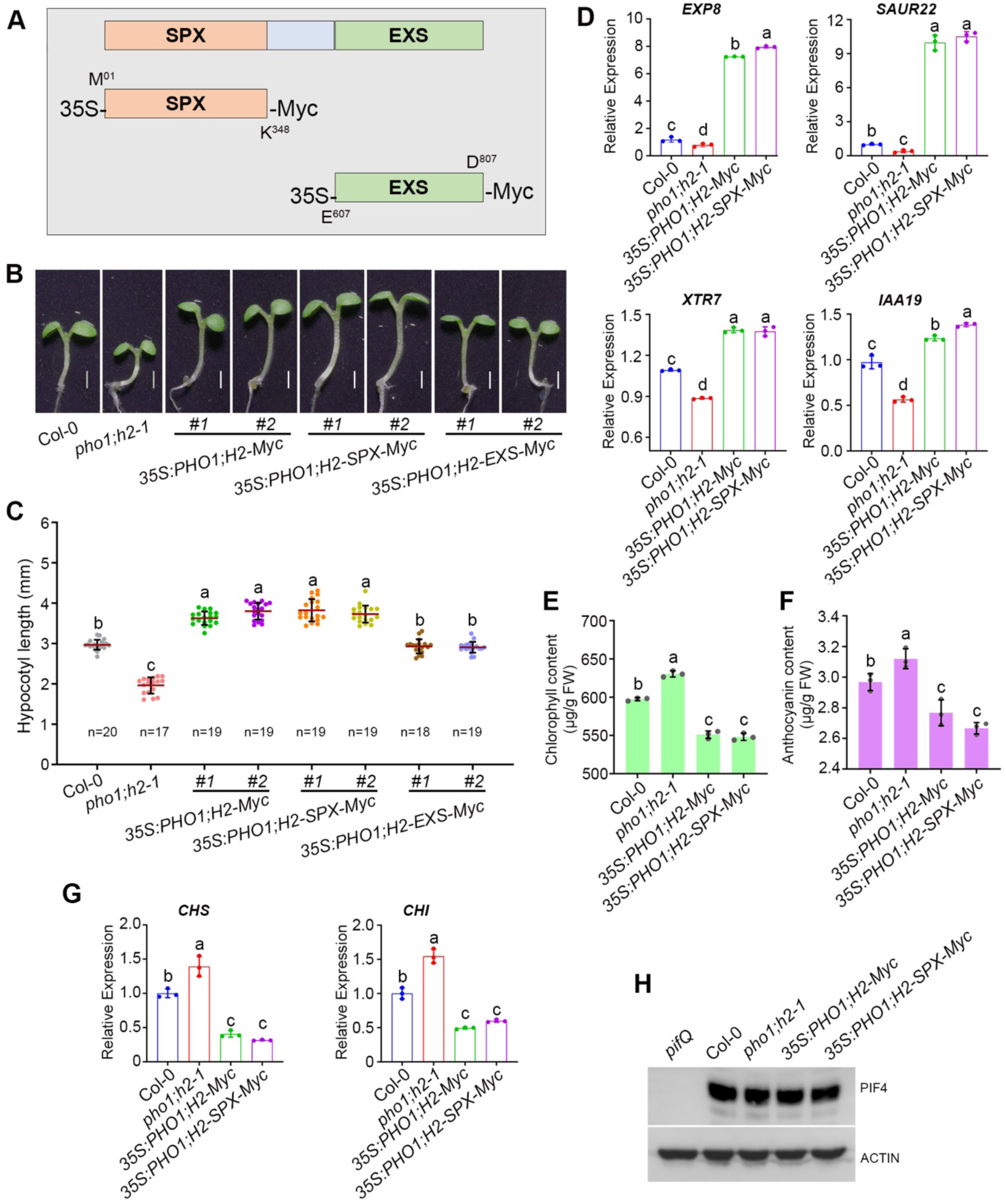
The SPX domain of PHO1;H2 is necessary and sufficient to suppress seedling photomorphogenesis in Arabidopsis. **(A)** Pictorial representation of full-length *35S:PHO1;H2-Myc* protein structure with *35s:PHO1;H2-SPX-Myc* (1-348 amino acids) and *35S:PHO1;H2-EXS-Myc* (607-807 amino acids). (B) Representative hypocotyl images of six-days-old seedling of Col-0*, pho1;h2-1, 35S:PHO1;H2-Myc #1, 35S:PHO1;H2-Myc #2, 35S:PHO1;H2-SPX-Myc #1, 35S:PHO1;H2-SPX-Myc #2, 35S:PHO1;H2-EXS-Myc #1* and *35S:PHO1;H2-EXS-Myc #2* grown in WL (60 µmol m²s¹) at 22°C under SD. The scale = 1 mm. (C) Hypocotyl length measurements of Col-0*, pho1;h2-1, 35S:PHO1;H2-Myc #1, 35S:PHO1;H2-Myc #2, 35S:PHO1;H2-SPX-Myc #1, 35S:PHO1;H2-SPX-Myc #2, 35S:PHO1;H2-EXS-Myc #1* and *35S:PHO1;H2-EXS-Myc #2* of seedlings shown in B. The error bars show standard deviation, and the red line marks the mean. n=indicates the number of seedlings measured. (**D-I**) RT-qPCR data reveal the transcript abundance of growth-promoting genes *EXP8* (D) *, IAA19* (E)*, XTR7* (F)and *SAUR22* (G), and light-inducing genes *CHS* (H) and *CHI* (I) in six-day-old seedlings grown in WL at 22°C under SD. *TUB2* was taken as an endogenous control. Relative gene expression was calculated by normalizing to Col-0 (set to 1). Bars denote the mean±SD (n=3 biological replicates). Seedling tissue harvested at ZT xx was used for the RNA extraction and RT-qPCR analysis. (**J** and **K**) Quantification of photopigments chlorophyll and anthocyanin content of six-day-old seedlings of Col-0*, pho1;h2-1, 35S:PHO1;H2-Myc , 35S:PHO1;H2-SPX-Myc genotypes* grown in WL (60 µmol m^-^²s^-^¹) at 22°C under SD conditions. Bars denote the mean±SD (n=3 biological replicates). (**G**) Immunoblotting with anti-PIF4 antibodies showing PIF4 protein levels in Col-0*, pho1;h2-1, 35S:PHO1;H2-Myc* and *35S:PHO1;H2-SPX-Myc* genotypes. Seedlings were grown in WL at 22°C under SD, and tissue was harvested at ZT4. The *pifQ* mutant was used as a negative control. Actin levels indicate equal protein loading. In C-K, the asterisk (*) represents the statistically significant differences from Col-0 and between the indicated pair of genotypes as revealed by Student’s *t*-test (\**p*<0.05; \*\**p*<0.001; ***p<0.001; ****p<0.0001). “ns” refers to non-significant.

Next, to determine whether the increased hypocotyl growth is due to misexpression of growth-promoting genes, we examined the expression levels of key growth-related genes, including *XTR7, IAA19, EXP8,* and *SAUR22* (27, 39–41). Consistent with the hypocotyl growth, these genes showed significant downregulation in the *pho1;h2-1* mutant compared to Col-0 (Fig. 5D). Interestingly, in both *PHO1;H2-Myc* and *PHO1;H2*-*SPX-Myc* overexpression lines, the expression of these genes was strongly upregulated compared to Col-0 (Fig. 5D), and their expression was largely comparable between *PHO1;H2-Myc* and *PHO1;H2*-*SPX-Myc* overexpression lines (Fig. 5D). Further, estimation of chlorophyll and anthocyanin contents revealed that they were significantly reduced in the *PHO1;H2-Myc* and *PHO1;H2*-*SPX-Myc* overexpression lines compared to Col-0 (Fig. 5E,F). However, the photopigment content was significantly elevated in the *pho1;h2-1* mutant compared to the Col-0 (Figure 5E,F). Consistent with the elongated hypocotyl growth phenotype and the reduced photopigment content, gene expression of light-inducible genes such as *CHS* and *CHI* that are involved in anthocyanin biosynthesis showed significant downregulation in *35S:PHO1;H2-Myc* and *35S:PHO1;H2*-*SPX-Myc* overexpression lines compared to Col-0. (Fig. 5G).

Given that many of the growth-responsive genes that are upregulated in the *PHO1;H2-Myc* and *PHO1;H2*-*SPX-Myc* overexpression lines, which are also direct transcriptional targets of PIF4 (27, 39–41), we were curious to know whether *PHO1;H2* or *PHO1;H2*-*SPX* overexpression influences the PIF4 protein levels. Immunoblotting with an anti-PIF4 antibody revealed that PIF4 protein abundance remained unchanged across all four genotypes, Col-0, *pho1;h2-1, PHO1;H2-Myc,* and *PHO1;H2*-*SPX-Myc* (Fig. 5H). These results suggest that either PHO1;H2 or the PHO1;H2-SPX domain does not affect the stability or accumulation of PIF4.

### PHO1;H2 levels affect the dynamic interaction between TOC1 and PIF4

Our findings reveal that PHO1;H2 physically interacts with TOC1 and negatively regulates photomorphogenesis, and also affects the expression of PIF4-target genes. However, the enhanced expression of PIF4 target genes is unlikely to be due to increased PIF4 protein abundance, as shown above (Fig. 5H). Earlier reports suggest that TOC1 inhibits PIF4 transcriptional activity, thereby reducing target gene expression (28). We thus hypothesize that PHO1;H2 enhances the transcriptional output of PIF4 target genes, potentially by counteracting TOC1 activity. To test this hypothesis, we performed *in vivo* and *in vitro* co-immunoprecipitation assays. For the *in vitro* competition assay, we expressed and purified recombinant His-TOC1 and GST-PIF4 from *E. coli*, and equimolar amounts of these proteins were incubated with total protein extracts from Col-0, *pho1;h2-1*, *35S:PHO1;H2-*Myc, and *35S:PHO1;H2-SPX-Myc* seedlings. We pulled down His-TOC1 by Ni-NTA resin, and the eluted pellet fraction was analyzed by immunoblotting using anti-GST and anti-His antibodies. The *pho1;h2-1* mutant showed higher levels of the immunoprecipitated PIF4 than the Col-0 (Fig. 6A). At the same time, in the *PHO1;H2-Myc* and *PHO1;H2-SPX-Myc* overexpression transgenic lines, the amount of PIF4 pulled down was visibly reduced compared to the Col-0 (Fig. 6A), indicating that elevated levels of PHO1;H2-FL or PHO1;H2-SPX in the overexpression lines effectively sequester TOC1 and reduce the TOC1 association with PIF4 (Fig. 6A).

**Fig. 6.**
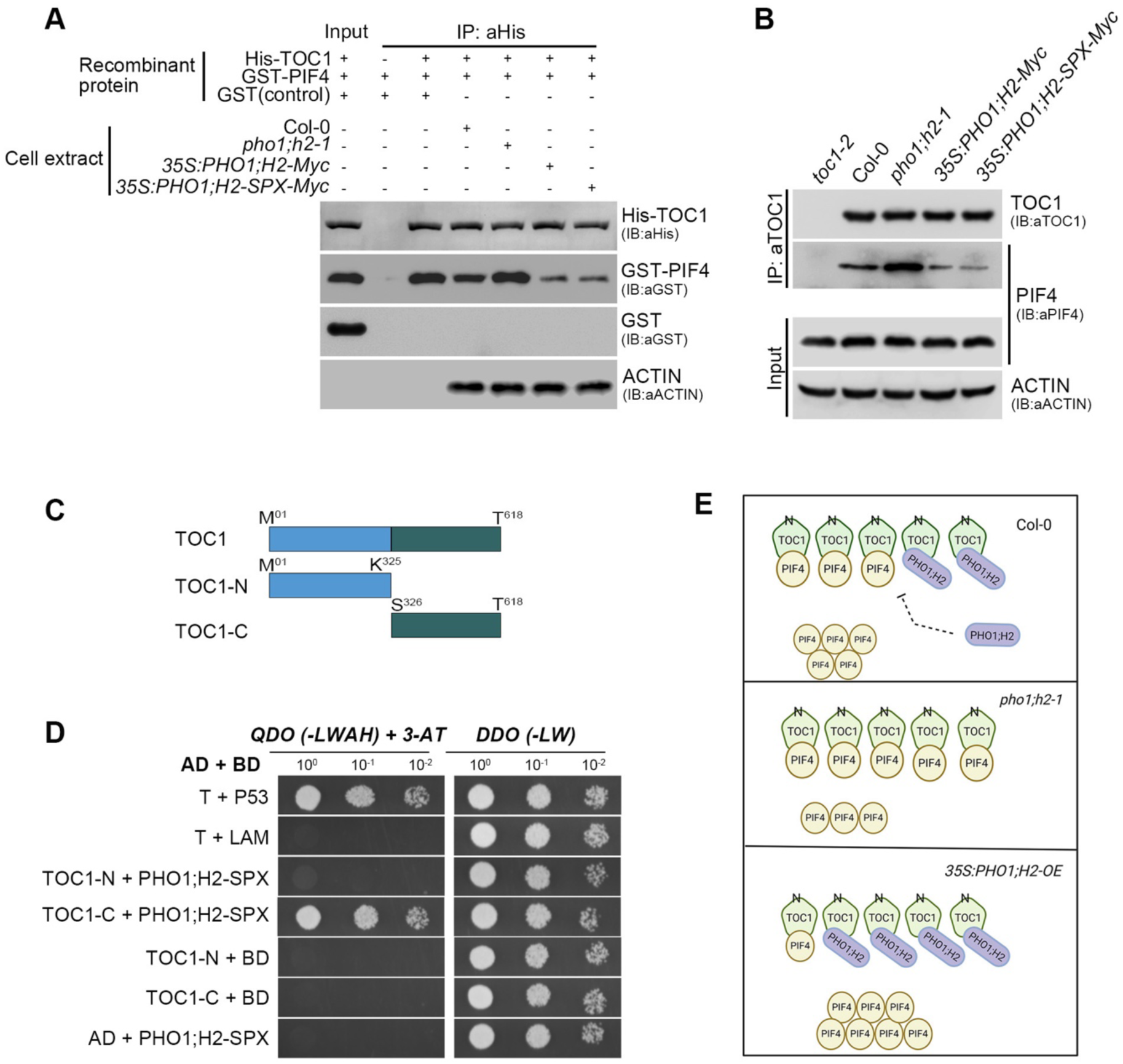
The SPX domain of PHO1;H2 physically interacts with the C-terminal region of TOC1, and likely interferes with its association with PIF4. (**A**) In vitro pull-down assay demonstrating co-immunoprecipitated GST-PIF4 with His-TOC1 incubated with the indicated cell extracts. Immunoblotting was done using anti-GST and anti-His antibodies. Acting was used as a reaction volume control. GST protein was used as a negative control. (**B**) In vivo co-immunoprecipitation of PIF4 along with TOC1 from Col-0, *pho1;h2-1*, *35S:PHO1;H2*, *35S:PHO1;H2-SPX* seedling extracts. The PIF4 protein amount, immunoprecipitated with TOC1, is higher in the *pho1;h2-1* mutant compared to the Col-0, while it is reduced in the *35S:PHO1;H2* and *35S:PHO1;H2-SPX* transgenic lines. (**C**) Schematic representation showing the domain architecture of TOC1. The full-length TOC1, the N-terminal part (blue) and the C-terminal part (Green) are shown based on annotated boundaries. (**D**) Yeast two-hybrid assay demonstrating that PHO1;H2-SPX physically interacts with TOC1 via the C-terminal domain. Co-transformants were first selected on double dropout medium (DDO, SD/-Leu/-Trp) to confirm the presence of both of bait and prey constructs (right panel), and subsequently cultures were dropped onto quadruple dropout (QDO, SD/-Ade/-His/-Leu/-Trp) medium to assess protein–protein interactions (left panel). (**E**) Schematic representation showing the competitive interaction between PHO1;H2, and PIF4 for binding to the C-terminal of TOC1. In Col-0, the *PHO1;H2* is insufficient to compete with PIF4, thereby leading to PIF4 sequestration by TOC1. In the *35S:PHO1;H2* (*PHO1;H2-OE*) line, elevated *PHO1;H2* levels result in stronger titration of TOC1 through binding to its C-terminal domain, preventing its association with PIF4 and maintaining a higher pool of PIF4.

Next, we carried out an *in vivo* Co-IP assay, pulling down TOC1 protein (using native TOC1 antibody) from Col-0, *pho1;h2-1, 35SPHO1;H2-Myc* and *35S:PHO1;H2-SPX-Myc* lines. In the co-immunoprecipitated samples, PIF4 was detected using an anti-PIF4 antibody. In the *pho1;h2-1* mutant the co-immunoprecipitated PIF4 protein was slightly more than Col-0 (Fig. 6B), whereas it was significantly reduced in the *35S:PHO1;H2-Myc* and *35S:PHO1;H2-SPX-Myc* transgenic lines (Fig. 6B), as revealed by the immunoblotting data, indicating that elevated *PHO1;H2-FL* or the *PHO1;H2-SPX* levels in the overexpression lines likely outcompete for TOC1 binding resulting in inhibition of TOC1-PIF4 association. (Fig. 6B). Together, these results confirm that PHO1;H2, via the SPX domain, binds TOC1 and likely inhibits its function through sequestration.

Previous reports suggest that TOC1 interacts with PIF4 via its C-terminal domain (28). To confirm whether PHO1;H2 also interacts with TOC1 through its C-terminal domain, we performed a yeast two-hybrid assay using the N-terminal SPX domain of PHO1;H2 and studied its interaction with the N-terminal and C-terminal domains of TOC1. We divided TOC1 into two fragments, N-terminal TOC1 (N-TOC1, 1-325 bp) and C-terminal TOC1 (C-TOC1, 326-618 bp) (Fig. 6C). N-TOC1 and C-TOC1 were fused to AD to generate N-TOC1-AD and C-TOC1-AD constructs, which were used as prey. The N-TOC1-AD and C-TOC1-AD constructs were separately co-transformed with PHO1;H2-SPX-BD. The Y2H assay results revealed that the SPX domain of PHO1;H2 specifically binds to the, C-terminal domain of TOC1 (C-TOC1, 326-618) as the yeast cells were able to grow on the synthetic QDO media containing 3-AT (Fig. 6D); however, the N-TOC1-AD failed to interact with PHO1;H2-SPX-BD, as the yeast cells could not grow on the synthetic QDO media containing 3-AT, suggesting that the C-terminal part of TOC1 is the common interaction site both for PIF4 and PHO1;H2 (Fig. 6D). Together, these multiple lines of evidence demonstrates that PHO1;H2, via its N-terminal SPX domain, binds to the C-terminal domain TOC1, where PIF4 also binds, and this binding is competitive, as evidenced by *in vivo* and *in vitro* pull-down assays and yeast two-hybrid results. As a result, PIF4 is released from TOC1-mediated repression when PHO1;H2 levels are higher (Fig. 6E).

### PHO1;H2 is essential for the optimal PIF4 function

The above results suggest that PHO1;H2 binding to TOC1 can suppress TOC1 binding to PIF4, alleviating TOC1-imposed suppression on PIF4 activity. To further substantiate these results at the genetic level, we crossed the *pho1;h2-1* mutant with *pif4-101* and generated the *pho1;h2-1 pif4-101* homozygous double mutant. We examined hypocotyl length in six-day-old seedlings under SD in WL. The *pho1;h2-1 pif4-101* double mutant exhibited hypocotyl length comparable to the *pif4-101* mutant, suggesting that the *pif4-101* mutation is epistatic to *pho1;h2-1* and PHO1;H2 is likely working in the same pathway as PIF4 (Fig. S7A,B).

Further, to check the effect of the *pho1;h2-1* mutation on PIF4-mediated hypocotyl growth, we introduced the *pho1;h2-1* mutation in the *PIF4-OE* background and generated a homozygous line and analysed hypocotyl phenotype in six-day-old seedlings grown in WL under SD photoperiod. *PIF4-OE* showed characteristic longer hypocotyls compared to Col-0, as reported (13) (Fig. 7A,B); however, in the presence of *pho1;h2-1*, the long hypocotyl phenotype of *PIF4-OE* was strongly suppressed in the *pho1;h2-1 PIF4-OE* background, as the hypocotyl length was much shorter than that of the *PIF4-OE* line (Fig. 7A,B), suggesting that PHO1;H2 is indeed required to promote PIF4 function and activity. Consistent with the mutant phenotype, overexpression of *PHO1;H2-Myc* and *PHO1;H2*-*SPX-Myc* in the *PIF4-OE* background resulted in further exaggerated hypocotyl length phenotype as seen in the *35S:PHO1;H2-Myc PIF4-OE* and *35S:PHO1;H2*-*SPX-Myc PIF4-OE* double homozygous transgenic lines, than the *PIF4-OE* line alone (Fig. 7A,B). Similarly, analysis of the hypocotyl length phenotype under LD photoperiod also suggests that the long hypocotyl phenotype of *PIF4-OE* is strongly suppressed by the *pho1;h2-1* mutant, while *PHO1;H2-Myc* or *PHO1;H2*-*SPX-Myc* overexpression further enhances hypocotyl growth (Fig. S7A,B).

**Fig. 7.**
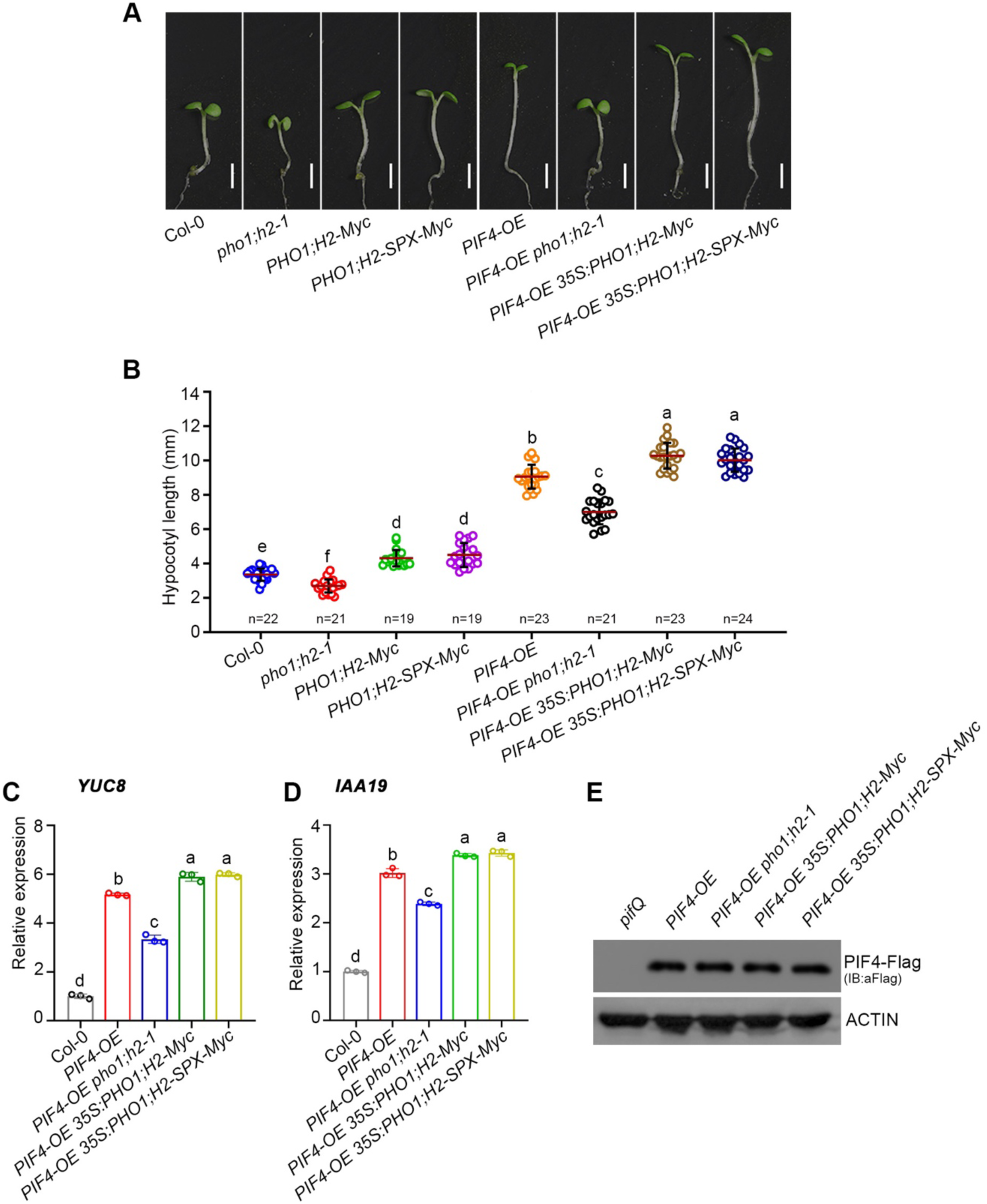
PHO1;H2 is essential for optimal PIF4 transcriptional activity and PIF4-mediated growth. (**A**). Representative hypocotyl images of six-day-old seedlings of Col-0*, pho1;h2-1, 35S:PHO1;H2-Myc , 35S:PHO1;H2-SPX-Myc , PIF4-OE, PIF4-OE pho1;h2-1, PIF4-OE 35S:PHO1;H2-Myc* and *PIF4-OE 35S:PHO1;H2-SPX-Myc* genotypes grown in WL (60 µmol m^-^²s^-^¹) at 22°C under SD photoperiod. Scale, 1 mm. (**B**) Hypocotyl length measurements of Col-0*, pho1;h2-1, 35S:PHO1;H2-Myc , 35S:PHO1;H2-SPX-Myc , PIF4-OE, PIF4-OE pho1;h2-1, PIF4-OE 35S:PHO1;H2-Myc* and *PIF4-OE 35S:PHO1;H2-SPX-Myc* genotypes shown in A. Error bars show standard deviation, and the red line marks the mean. “*n”* indicates the number of seedlings measured. (**C** and **D**) RT-qPCR analysis of PIF4 downstream of genes *YUC8* (C) and *IAA19* (D) in Col-0*, PIF4-OE, PIF4-OE pho1;h2-1, PIF4-OE 35S:PHO1;H2-Myc* and *PIF4-OE 35S:PHO1;H2-SPX-Myc* genotypes. The six-day-old seedlings grown in WL at 22°C under SD were used for the RT-qPCR analysis. *TUB2* was taken as an endogenous control. (**E**) Endogenous PIF4 protein levels in*, PIF4-OE, PIF4-OE pho1;h2-1, PIF4-OE 35S:PHO1;H2-Myc* and *PIF4-OE 35S:PHO1;H2-SPX-Myc* lines. The *pifQ* mutant was used as a negative control. Actin protein levels serve as a loading control. In Figs. B-D, asterisks represent the statistically significant differences from Col-0, or between the indicated pair of genotypes as revealed by Student’s *t*-test (***p<0.001; ****p<0.0001). “ns” refers to non-significant.

To further validate the PHO1;H2-, or N-terminal SPX-domain-mediated enhancement of PIF4 activity, we measured transcript abundance of key PIF4 target genes, such as *YUC8* and *IAA19,* by RT-qPCR. As observed previously (13), the expression of *YUC8* and *IAA19* was significantly upregulated in the *PIF4-OE* line compared with Col-0 (Fig. 7C,D). However, in the *pho1;h2-1 PIF4-OE* background, the expression of *YUC8* and *IAA19* genes showed significant downregulation compared to *PIF4-OE* (Fig. 7C,D). In contrast, in the *35S:PHO1;H2-Myc PIF4-OE, 35S:PHO1;H2-SPX-Myc PIF4-OE* double transgenic lines, *YUC8* and *IAA19* gene expression was significantly enhanced than the *PIF4-OE* line (Fig. 7C,D), Moreover, examining the PIF4 protein levels in the *PIF4-OE*, *pho1;h2-1 PIF4-OE, 35S:PHO1;H2-Myc PIF4-OE, 35S:PHO1;H2-SPX-Myc PIF4-OE* lines reveal that the PIF4 protein levels are unaltered in these lines and which is comparable to the *PIF4-O*E line (Fig. 7E), suggesting that either absence or elevated levels of PHO1;H2 do not affect the PIF4 protein levels. Also, our Y2H assay revealed that PHO1;H2 do not physically interact with PIF4 (Fig. S8). These data confirm that the observed upregulation of PIF4 target gene expression is attributable to enhanced PIF4 transcriptional activity, driven by increased protein abundance of PHO1;H2 or PHO1;H2-SPX. Collectively, our above genetic, molecular, and biochemical data support our hypothesis that PHO1;H2, through its SPX domain, binds to TOC1 and sequesters it, relieving repression of PIF4 and enhancing expression of auxin signaling and growth-responsive genes (Fig. 8).

**Fig. 8.**
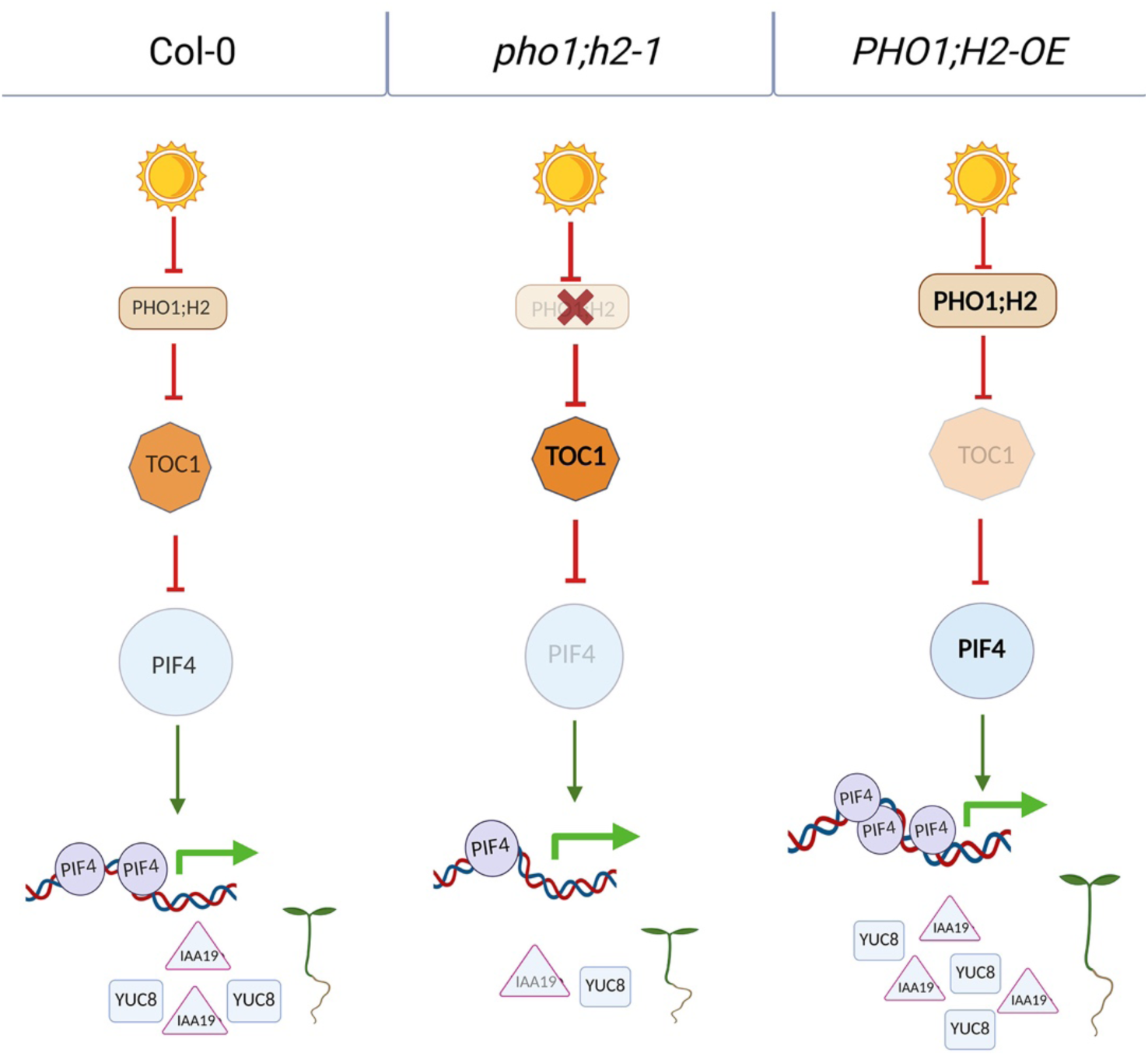
Hypothetical model demonstrating the molecular mechanism of PHO1;H2 function. A mechanistic model describing the role of PHO1;H2 in regulating seedling photomorphogenesis. Under wild-type (Col-0) conditions, moderate PHO1;H2 levels maintain optimal photomorphogenic growth. TOC1 remains partially active in the wild-type, in which a fraction of PHO1;H2 binds TOC1, while the remaining TOC1 interacts with PIF4, sequestering it, leading to optimal photomorphogenic growth. Although this reduces the pool of free PIF4, some remains available to bind to the promoters of growth-associated genes such as *IAA19* and *YUC8*. In the *pho1;h2-1* mutant background, the absence of *PHO1;H2* removes this regulatory checkpoint, resulting in enhanced TOC1 activity. Consequently, PIF4 is fully sequestered by TOC1, preventing its association with promoters of downstream growth-promoting genes. By contrast, in the *PHO1;H2-OE* line, the elevated levels of PHO1;H2, particularly its N-terminal SPX domain, bind to the C-terminal part of TOC1 and strongly inhibit its association with PIF4. This competitive interaction relieves PIF4 from TOC1-mediated sequestration, allowing it to activate downstream target genes such as *IAA19* and *YUC8*, thereby enhancing hypocotyl growth and dampening photomorphogenic responses.

## Discussion

Light and the circadian clock play critical roles in the spatio-temporal regulation of gene expression, orchestrating plant growth and development (45, 47). Light not only entrains the circadian oscillator through photoreceptor-mediated signaling but also gates clock output to align physiological responses with the external photoperiod (48). Despite this, the molecular intricacies of the dynamic regulation between light and clock components are poorly understood. This study uncovered that PHO;H2 acts as a novel repressor of seedling photomorphogenesis, functioning downstream of all the light wavelengths tested (Fig. 1). Consistent with the reduced hypocotyl length observed in the *pho1;h2-1* mutant, the expression of various growth-promoting genes such as *YUC8, IAA19, SAUR22, EXP8* and *XTR7* is significantly downregulated.

However, the light-induced genes *CHS, CHI, CAB1, RBCS1A,* and *ELIP2* showed significant upregulation (Fig. 2), suggesting that PHO1;H2 differentially regulates the expression of genes involved in hypocotyl growth and de-etiolation. Further, our study demonstrates that PHO1;H2 physically interacts with TOC1 through its N-terminal SPX domain and likely suppresses its function as revealed through epistatic analysis (Figs. 3,4). This mechanistic insight is supported by functional assays using domain-specific overexpression constructs. Overexpression of full-length PHO1;H2 and the PHO1;H2-SPX domain alone led to significant hypocotyl elongation, whereas overexpression of the EXS domain alone failed to induce such growth, indicating that the SPX domain is both necessary and sufficient for this regulatory function (Fig. 5). This supports our conclusions that the physical interaction between the SPX domain and TOC1 is a key determinant in modulating PIF4 activity.

Furthermore, gene expression analysis of PIF4 target genes provides strong evidence of this regulatory relationship. In the *pho1;h2-1* mutant, which lacks functional PHO1;H2, canonical PIF4 target genes such as *EXP8, SAUR22, XTR7* and *IAA19* are significantly downregulated while the light-induced genes such as *CHS* and *CHI* showed significant upregulation, consistent with reduced hypocotyl growth but enhanced photomorphogenic responses, which is likely due to TOC1-mediated suppression of PIF4 activity. Conversely, these same target PIF4 target genes are upregulated in both *35S:PHO1;H2* and *35S:PHO1;H2-SPX* lines (Fig. 5D-E), indicating that the primary function of PHO1;H2 is to relieve TOC1-imposed repression on PIF4, thereby enabling PIF4 to activate transcription of genes required for cell elongation and hypocotyl growth. Accordingly, elevated PHO1;H2 attenuates TOC1 activity, reducing expression of TOC1-dependent photomorphogenic genes. Notably, TOC1 is known to activate *CAB* expression and promote photomorphogenesis (45). Therefore, the enhanced expression of *CAB1* and other photomorphogenesis-related genes in the *pho1;h2-1* is due to enhanced TOC1 activity, while reduced expression of these genes in the *35S:PHO1;H2* and *35S:PHO1;H2-SPX* lines is due to reduced TOC1 activity.

TOC1 has previously been shown to interact with PIF4 via its C-terminal domain, thereby repressing PIF4-mediated transcription of growth-promoting genes (28). We now show that the SPX domain of PHO1;H2 binds to the same C-terminal region of TOC1 responsible for PIF4 interaction (Fig. 6), suggesting a competitive binding mechanism in which PHO1;H2 displaces TOC1 from PIF4, liberating a transcriptionally competent pool of PIF4 to drive hypocotyl elongation. This places PHO1;H2 in a regulatory layer distinct from, yet complementary to, the CCA1–SHB1 module, which promotes *PIF4* transcription during early light phases by direct binding to its promoter (29). While CCA1–SHB1 controls PIF4 transcript accumulation, PHO1;H2 modulates PIF4 activity post-translationally by antagonizing TOC1 through competitive binding (Figs. 3–6). This dual-layered regulation ensures that *PIF4* is not only transcribed at the appropriate time, but also that its transcriptional activity is appropriately modulated by the circadian clock in response to dynamic light cues.

The PHO1 family proteins have been traditionally characterized for their role in phosphate loading and transport (36, 38, 49, 50). Identifying PHO1;H2 as an inhibitor of photomorphogenesis via PIF4 signaling expands the functional repertoire of this family. Given the widespread presence of SPX domains in phosphate-responsive proteins and their established role in sensing inositol pyrophosphate signals (34, 50–54), it is plausible that PHO1;H2-mediated interference with TOC1 could itself be dynamically regulated by phosphate availability or metabolic status, although this remains to be tested.

Collectively, this study establishes PHO1;H2 as a negative regulator of photomorphogenesis by modulating the TOC1–PIF4 interactions (Fig. 8). The SPX domain of PHO1;H2 emerges as a crucial effector of this activity, enabling PHO1;H2 to control hypocotyl elongation by selectively releasing PIF4 from TOC1-imposed inhibition (Fig. 8) These results emphasize the complexity of the transcriptional and post-translational circuits that orchestrate plant growth responses through coordinated function between light and the circadian clock signaling pathways.

## Materials and Methods

### Plant materials and growth conditions

For this study, the wild-type ecotype Columbia (Col-0) of *Arabidopsis [(Arabidopsis thaliana (L.)]* was used for all experiments. The *PHOSPHATE 1 HOMOLOG 2* (*PHO1;H2*) mutants *pho1;h2-1* (SALK_127006) and *pho1;h2-2* (GK-337F03.04) were obtained from Eurasian Arabidopsis Stock Centre (euNASC). The *mutants* (*toc1-2)* (*45*)*, pif4-102* (55) and the transgenic line *PIF4-OE* (*pPIF4:PIF4:FLAG)* were previously described (56). The *phyb-9*, *cop1-4*, *hy5-215,* and *elf3-4* were described elsewhere (57–60).

### Identification of T-DNA insertion mutants in the *PHO1;H2* locus

All the genotyping samples were grown on soil, and then a 2x2 mm leaf disc was cut and dipped into 0.25 M sodium hydroxide and boiled for 30 seconds at 100°C, followed by adding 0.25 M hydrochloric acid (HCl) and 0.5 M Tris-HCl buffer (pH 8.0) and again boiled for 2 min at 100°C. The samples were then used as templates. For T-DNA verification, PCR was performed using the T-DNA Border primer and a gene-specific right primer. The T-DNA lines were further confirmed by RT-PCR. The primer list is in Table S1.

### Generation of *PHO1:H2 Myc* overexpression transgenic lines

To generate the full-length PHO1;H2 overexpression lines, the full-length CDS of the *PHO1; H2* gene (2424 bp) was amplified from WT Col-0 cDNA. Using Gateway technology, the amplified PHO1 H2 CDS was cloned into a pENTR^TM^ Directional TOPO^R^ vector and confirmed by PCR. The PHO1; H2 pENTR (Invitrogen) construct was then cloned into the destination vector pGWB617 vector 4X-Myc epitope tag at C-terminus *(att*R1-Cm^r^-*ccd*B-*att*R2-10xMyc-T_NOS_) (61) by gateway-based LR recombination. The Col-0 plants were then transformed using the *Agrobacterium*-mediated floral dip (62). The obtained T_0_ plants were plated, and T_1_ plants were selected on phosphinothricin. Several independent transgenic lines were transferred to soil and confirmed by genotyping using PCR with a 35S promoter primer and a gene-specific primer. Finally, in the T_3_ generation, two independent homozygous transgenic lines were used for further phenotypic and molecular analysis.

### Generation of higher-order mutants for genetic studies

To generate the double mutant *pho1;h2-1 toc1-2, pho1;h2-1* was crossed with the *toc1-2* mutant, and F_2_ seeds were plated on MS media, and double mutants *pho1; h2-1 toc1-2* were selected based on phenotypic traits and were confirmed for the genotype using PCR. To make *pho1;h2-1 pif4-101* and *pho1;h2-1x35s:PIF4OE* lines, the *pho1; h2-1* mutant was crossed with *pif4-101* and *PIF4 OE.* The F_1_ seeds were planted to get F*2* seeds. The F_2_ seeds were plated on phosphinothricin and kanamycin, respectively, to select *pho1;h2-1 pif4-101* and *pho1;h2-1 PIF4-OE* double mutant combinations. Homozygosity was confirmed by PCR-based genotyping.

### Generation of *PHO1;H2-SPX-Myc* and *PHO1;H2-EXS-Myc* overexpression transgenic lines

SPX and EXS domains were amplified from the *PHO1;H2:Myc* construct. Amplified SPX and EXS domains were cloned into pENTR^TM^ Directional TOPO^R.^ Both the SPX and EXS entry vectors were cloned into the destination vector pGWB420 *(att*R1-Cm^r^-*ccd*B-*att*R2-10xMyc-T_NOS_) using LR Clonase. These binary vectors were introduced into the *Agrobacterium tumefaciens,* and the wild-type Col-0 was transformed using the floral dip method. T_0_ seeds were selected based on relevant antibiotic resistance. PCR confirmation using 35S primer. All experiments were conducted using homozygous seeds from the T_3_ and T_4_ generations. Primers used are listed in Table S1.

### Generation of PHO1;H2-Myc PIF4-OE and PHO1;H2-SPX-Myc PIF4-OE double transgenic lines

To generate *PHO1;H2-Myc PIF4-OE* and *PHO1;H2-SPX-Myc PIF4-OE* double transgenic lines, the *PHO1;H2 Myc* and *PHO1;H2-SPX-Myc* lines were crossed to *PIF4-OE* transgenic line. The F_1_ seeds were harvested and planted to get F_2_ seeds. The F_2_ seeds were plated on kanamycin and basta, and the seedlings containing double transgenes were identified and transferred to soil. In the F_3,_ putative double-homozygous lines were identified based on phenotype and antibiotic selection and further confirmed by PCR in the adult stage. F_4_ seeds were harvested, confirmed by antibiotic selection and phenotype, and used for the other assays.

### Chlorophyll and anthocyanin measurements

Chlorophyll and anthocyanin contents were estimated essentially as described by (21) and (22), with slight modifications. Six-day-old seedlings were harvested in an Eppendorf and snap-frozen in liquid nitrogen. About 30 mg of seedlings were ground into a powder, and 1 mL of chilled 80% acetone was added. The lysate was centrifuged for 10 min. at 20,000g. After centrifugation, the supernatant was transferred to a fresh Eppendorf tube, and the OD_645_ and OD_663_ were measured for chlorophyll a and chlorophyll b, respectively. Chlorophyll contents, expressed as ChlA=12.7*A_645_-2.69*A_663_ and ChlB=22.9*A_645_-4.48*A_663_ per gram of fresh seedlings, were calculated.

For anthocyanin quantification, 30 mg of seedlings were harvested and crushed in liquid nitrogen. The extract was prepared by adding 400 μL of extraction buffer (1% HCl in methanol). The samples were vortexed and kept at 4°C in the dark overnight. 200 μl of sterile H_2_O and 200 μl of CHCl_3_ were added, and the mixture was centrifuged at 13,000 rpm for 10 minutes. The supernatant was collected in a microcentrifuge tube. Anthocyanin content was expressed as (A_530_-0.33*A_657_) per gram of fresh seedlings.

### Measurement of hypocotyl length

The seeds were surface-sterilized with a sterilization solution (70% ethanol +0.5% Triton X-100), followed by washing with 100% ethanol (Merck EMSURE), and then dried on autoclaved filter paper in the laminar hood. The surface-sterilized seeds were plated on MS media (Sigma, M5519) and kept at 4°C for four days in the dark for stratification. After four days, the plates were moved to 22°C LD (16h-light/8h-dark) chamber for germination for a day. The plates were then transferred to short-day conditions at 22°C (8 h light/16 h dark) with different wavelengths of light at a fixed fluence rate. Six-day-old seedlings were aligned, and hypocotyls were measured using ImageJ (http://rsbweb.nih.gov/ij/). Statistical analysis was performed using GraphPad Prism.

### Yeast two-hybrid (Y2H) assay

To detect the interaction between *PHO1;H2*, and *TOC1*, we generated yeast two-hybrid clones. The *PHO1;H2* (2424bp) was amplified and digested with *Xma1* and *BamH1*, followed by ligation into the vector pGBKT7*-BD* (containing the *GAL4* DNA binding-domain) (Takara Bio, Shiga, Japan) as bait with the intact reading frame. The TOC1 (1857 bp) was amplified and digested with *EcoRI* and *XmaI* and ligated into pGBKT7 (Takara Bio) containing the *GAL4* Activation domain (AD) sequence to serve as prey. Both PHO1;H2-BD and TOC1-AD were co-transformed into yeast strain Y2H GOLD and plated on double dropout media (SD /-W/-L). The colonies were then tested for growth on the quadruple dropout media (SD -W/-L/-A/-H) plates to investigate the interactions.

For domain-specific interaction studies, N-terminal TOC1 (975 bp) and C-terminal TOC1 (879 bp) were amplified with primers given in Table S1. The amplified product was digested and ligated into the GAL4 activation domain. The nTOC1-AD and cTOC1-AD in combination with SPX-BD were co-transformed into yeast. Colonies from double dropout plates were screened on quadruple dropout plates. Primers used in the study are listed in Table S1.

### Bimolecular fluorescence complementation (BIFC) assay

Full-length TOC1 was amplified and cloned into pENTR^TM^ Directional TOPO^R^. The pENTR clone was then used to introduce the TOC1 gene into the 778 *pE-SPYNE* destination vector to get *TOC1-nYFP*. Similarly, *PHO1;H2* entry clone was used to introduce *PHO1;H2* into the 777-*pE-SPYCE* vector to get *PHO1;H2-cYFP*. For domain-specific BIFC constructs, *SPX-cYFP* and *EXS-nYFP* clones were generated using LR reactions. Each of these constructs was then transformed into the *Agrobacterium tumefaciens* GV3101 strain. Combinations of *Agrobacterium* (containing desired constructs; PHO1;H2-cYFP + TOC1-nYFP, SPX-cYFP+TOC1-nYFP, EXS-cYFP+TOC1-nYFP) were co-infiltrated into onion epidermal cells along with *p19* construct (*63*). For positive and negative controls, we used LRB1-nYFP + LRB2-cYFP and empty vectors nYFP + cYFP, respectively. Images were taken using an Olympus IX81 epifluorescence microscope. DAPI (4’,6-diamidino-2-phenylindole) was used to mark the nuclei. Primers used in the study are listed in Table S1.

### Total protein extraction and immunoblotting

Six-day-old seedlings, grown on MS medium, were harvested into liquid nitrogen and crushed. Total protein was extracted with protein extraction buffer containing (150mM NaCl, 50mM Tris-HCl at pH 8, 0.5M EDTA, 0.01% NP40, 5% glycerol, 5 μM phenylmethylsulfonyl fluoride (PMSF), 20 μM ß-mercaptoethanol, 20 μM DTT, 1X protease inhibitor cocktail (Roche). Samples were centrifuged at 13000 rpm at 4°C for 20 min, and the supernatant was collected. Protein was quantified using Bradford’s assay. Equal amounts of protein were loaded and separated on 10% (w/v) SDS-PAGE and transferred to PVDF membrane (Millipore IMMOBILON Merck). The gel blot was analyzed using the SuperSignal West Pico Plus substrate kit (Thermo Scientific, Waltham, USA). PIF4 was detected using goat polyclonal anti-PIF4 antibody (Agrisera), ACTIN was detected using rabbit polyclonal anti-ACTIN antibody (Agrisera), and TOC1 was detected using rabbit polyclonal anti-TOC1 antibody (PhytoAB).

### In-vitro pull-down of SPX-GST and TOC1-HIS

The SPX domain coding sequence was amplified and cloned into the pGEX4T2 vector to generate the GST-SPX construct. Similarly, TOC1 was cloned into the pET28a vector to get His-TOC1. *E. coli* BL21 was transformed with these vectors and used for protein production. Proteins were expressed and purified from *E. col*i after inducing with 0.1mM of IPTG for 2 hours at 37°C. GST-SPX was purified by Glutathione Sepharose Beads (Gold Bio). His-TOC1 was pulled by Ni-NTA Agarose Beads (Qiagen). Equimolar amounts of GST or GST-SPX were incubated with His-TOC1 bound to 50 μL beads in parallel reactions overnight at 4°C. The beads were washed 5 times with washing buffer (50 mM Tris-Cl, pH 8, 150 mM NaCl, 5% [w/v] glycerol, 1% ß-mercaptoethanol). The beads were then collected in fresh tubes and boiled with the sample buffer (50 mM Tris-Cl, pH 8, 150 mM NaCl, 15 % [w/v] glycerol, 4 % SDS, 20 mM DTT) for 10min at 70°C before separating on SDS-PAGE. Rabbit raised polyclonal anti-GST and anti-His antibodies were used to detect GST-SPX and His-TOC1, respectively. GST served as the negative control.

### In-vitro pull down assay to check the competition between PIF4-GST and TOC1-HIS, and TOC1-HIS with PHO1;H2-Myc *or* PHO1;H2-SPX-Myc

Full-length PIF4 was amplified and cloned into the pGEX4T2 vector. Recombinant GST-PIF4 was expressed and purified from *E.coli* BL21(DE3). GST-PIF4 was purified by Glutathione Sepharose beads (Gold Bio., St Lois, MI, USA). The TOC1-HIS protein was purified by Ni-NTA agarose beads (Qiagen). For the preparation of total protein extract, six-day-old seedlings of Col-0*, pho1;h2-1, PHO1;-H2-Myc and PHO1;H2-SPX-Myc* were harvested and crushed in liquid nitrogen. Plant extracts were prepared in the pull-down buffer (50 mM Tris-Cl, pH 8, 150 mM NaCl, 5% [w/v] glycerol, 1 mM EDTA, 1 mM phenylmethylsulfonyl fluoride (PMSF), 0.1% [v/v] Nonidet P-40, and 2X protease inhibitors cocktail; Roche) with 50 μM MG132. Samples were centrifuged at 20,000g for 20 min at 4°C, and the supernatant was collected into new vials.

To assess the effect of PHO1;H2 or PHO1;H2-SPX on the TOC1-PIF4 interaction, Gst-PIF4 and His-TOC1 (1.5 μg each) proteins were incubated with 250 μg of plant extracts of Col-0, *pho1;h2-*1, *PHO1;H2-Myc and PHO1;H2-SPX-Myc* for 2 hr at room temperature. His-TOC1 was pulled down as prey using 50 μL of Ni-NTA agarose beads by centrifugation at 2000 rpm, and the supernatant was removed. Excess unbound protein was washed off by repeating step thrice with 1ml of wash buffer (50 mM Tris-Cl, pH 8, 150 mM NaCl, 0.2% [v/v] glycerol, 1% ß-Mercaptoethanol). The pellet was resuspended in 25 μL of pull-down buffer and 5 μL of 6x loading dye, then boiled for 5 min and analyzed by SDS-PAGE. Rabbit polyclonal antibodies against the His and GST tags were used for immunodetection.

### *In-vivo* competition assay using transgenic and mutant genotypes

Six-day-old seedlings of Col-0, *pho1;h2-1, PHO1;H2-Myc and PHO1;H2-SPX-Myc* were harvested and crushed in liquid nitrogen. Total proteins were extracted in CO-IP buffer (50 mM Tris-Cl, pH 8, 150 mM NaCl, 5% [w/v] glycerol, 1 mM EDTA, 1 mM phenylmethylsulfonyl fluoride (PMSF), 0.1% [v/v] Nonidet P-40, and 2X protease inhibitors cocktail; Roche). Samples were centrifuged at 20,000g, and the supernatant was collected.

Co-immunoprecipitation was performed using Dynabeads^TM^ (Invitrogen) according to the manufacturer’s protocol, with some modifications. Briefly, 20 μL of beads were incubated with 3 μg of anti-His antibody for 2 hours at room temperature. Beads were washed twice with wash buffer (Invitrogen) and incubated with 300 μg of protein extract overnight. Beads were washed thrice and collected into new vials. Beads were then boiled with 30 μL of elution buffer with added Laemmli buffer for 10 min at 70°C. Elute was then separated on SDS-PAGE. An anti-PIF4 antibody was used to detect PIF4 levels.

### Phylogenetic analysis

Coding sequence for *PHO1;H2* was retrieved from TAIR. The FASTA sequence was then aligned using NCBI BLAST to identify homologs across different plant genomes. Homologs from different species were retrieved in FASTA and uploaded to NCBI COBALT for the alignment. MEGA 11 (Molecular Evolutionary Genetics Analysis Version 11) and the LG MODEL was used to generate the phylogenetic tree.

### Structure modelling and molecular docking for protein-protein interactions

The coding sequences of *PHO1;H2*, N-terminal SPX, and C-terminal EXS were submitted to I-TASSER to obtain the 3D structure. The 3D structure was visualized in pYMOL. The modelled structures of PHO1;H2-SPX, and TOC1 were used for molecular docking in CLUS PRO. The generated structure was then analysed, and the TOP-rank model was chosen for further analysis. For analyzing the TOC1-bound SPX complex and the residual interaction between SPX and TOC1, the structure was submitted to PDBsum. PDBsum provides a list of amino acids that interact with or form the interface of each protein. Amino acids were then marked in the structure using PYMOL.

### RNA Isolation, cDNA synthesis and RT-qPCR analysis

The RNeasy Plant Mini Kit (Qiagen) was used for RNA extraction according to the manufacturer’s protocol. Six-day-old seedlings were harvested, crushed in liquid nitrogen, and vortexed with 450 μL of buffer RLC. Lysate was transferred to the QIAshredder spin column and centrifuged for 2 min at 20,000g. The flow-through supernatant was transferred to a new vial, and 500 μL of ethanol was added. The solution was then transferred to the RNeasy Mini spin column and spun for 30 sec at 10,000 g. The column was washed with 700 μL of buffer RW1, followed by 500 μL of buffer RPE, and centrifuged at 10,000g for 2 min. The membrane was dried by centrifuging again for 1 min at 10,000g. The RNA was eluted by adding 50 μL of RNase-free water.

For cDNA preparation, Superscript III reverse transcriptase (Invitrogen) was used, and 1.5 μg of RNA was converted into cDNA. cDNA was diluted in a 1:20 ratio, and 3 μL of cDNA was used for one reaction. qPCR was carried out in QuantStudio5, Applied Biosystems, Thermo Fisher Scientific, using Power Up^TM^ SYBR Green Master Mix (Applied Biosystems). *TUBULIN β-2* (AT5G62690) was used as an endogenous control. Primers used in this study are listed in Table S1.

### Quantitative and statistical analyses

ImageJ (https://imagej.nih.gov/) was used to measure hypocotyl length. Students’ t-test and ANOVA followed by Tukey’s post-hoc HSD test (p<0.05) were used to determine significance between genotypes or time points. GraphPad Prism was used to make graphs and statistical analyses.

### Accession numbers

Information about the genes used in this study can be found in The Arabidopsis Information Resource database under the following accession numbers:, *AT2G03260 (PHO1;H2), PIF4 (AT2G43010), CCA1 (AT2G46830), ELF3 (AT2G25930), ELF4 (AT2G40080), PRR5 (AT5G24470), PRR7(AT5G02810), TOC1 (AT5G61380), AT1G29930 (CAB1), AT5G13930 (CHS), AT2G43570 (CHI), AT1G67090 (RBCS1A), AT4G14130 (XTR7), AT2G40610 (EXP8), AT5G62690 (TUB2), AT2G37620 (ACTIN1), YUC8 (AT4G28720), ELIP2 (AT3G22840), IAA19 (AT3G15540), SAUR22 (AT1G29430), RBCS1A (AT1G67090)*.

## Data and materials availability

All other data presented in the manuscript are available in the main manuscript and in the supporting information.

### Author contributions

S.N.G. conceptualized the idea, acquired funding, designed experiments, supervised the study, and edited the final drafts of the manuscript with inputs from D.D. and C.S. D.D. conceptualized and designed the experiments, generated the necessary reagents, performed most experiments, collected results, analyzed the data, prepared figures and wrote the first draft of the manuscript. C.S. conceptualized and carried out key experiments related to protein-protein interaction studies and also contributed to preparing the first draft. B.C.M. contributed to the initial screening and identification of the *pho1;h2* mutant. All the authors have read and approved the final version of the manuscript.

## Acknowledgements

The *TOC1* mutant (*toc1-2)* was kindly gifted by Dr Manjul Singh from CSIR-NBRI, India. D.D. acknowledges IISER Kolkata (Ministry of Education) for the doctoral fellowship. C.S. and B.C.M. acknowledge the Council of Scientific and Industrial Research (CSIR) for their doctoral fellowships. We thank all the members of Gangappa’s laboratory for fruitful discussions.

## Supporting Information

### Supplemental data includes

**Fig. S1.**
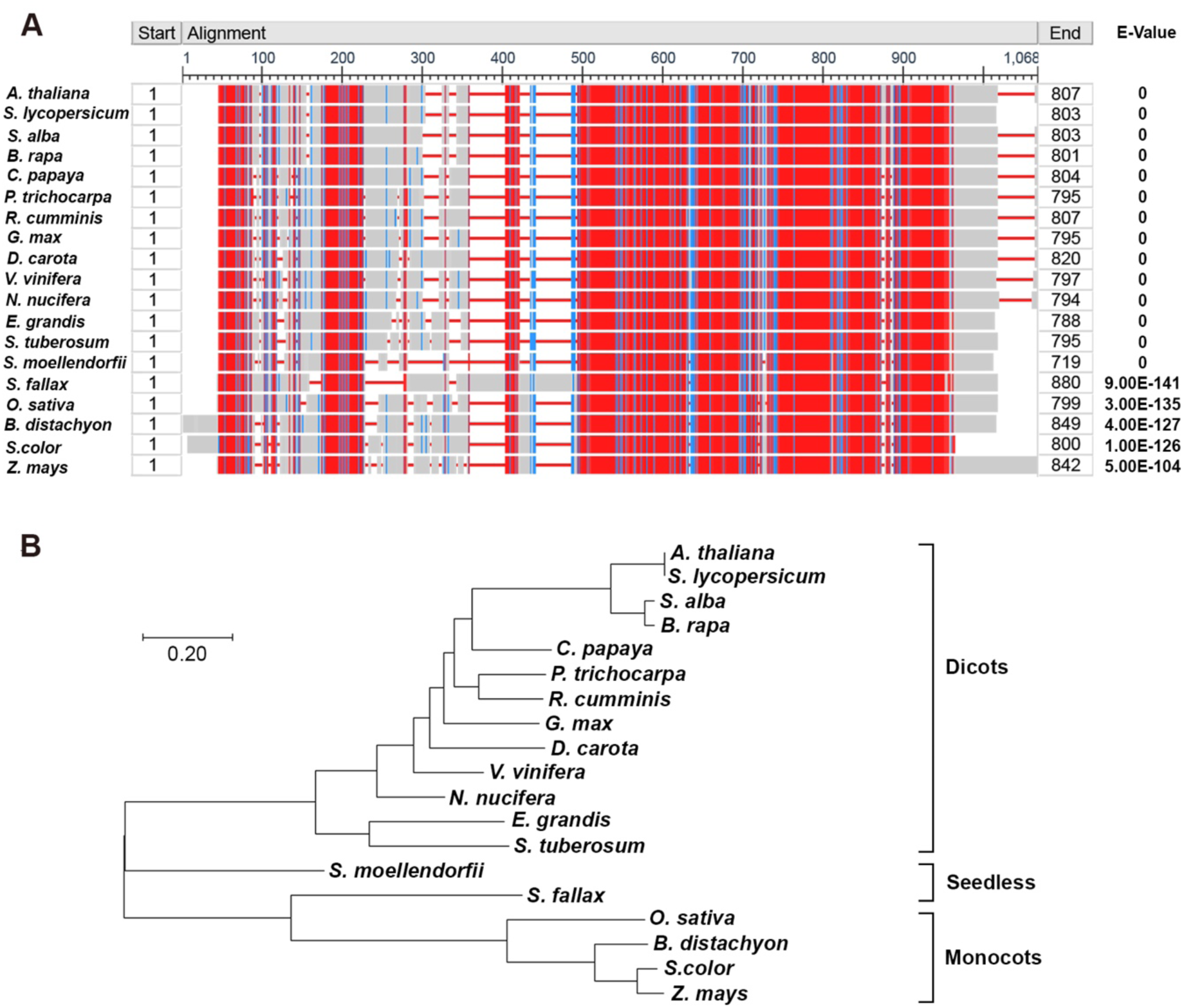
PHO1;H2 is a homolog of the PHO1 family of proteins, and is highly conserved across plant species. (**A**) Phylogenetic tree of PHO1 HOMOLOG 2 protein across different plant genomes of Dicots, monocots and seedless plants. The tree is drawn to scale with a branch length of 0.2. Based on 1,000 iterations, the bootstrapping values are shown below the branches. (**B**) Protein sequence alignment of PHO1;H2 identified by BLASTp against the different plant genomes on NCBI with calculated E-values. The identified Protein sequence was aligned with NCBI COBALT, with red indicating highly conserved and blue indicating low-conserved regions. The alignment results highlight domains in red, indicating high conservation and sequence similarity, while blue indicates low conservation or variable regions.

**Fig. S2.**
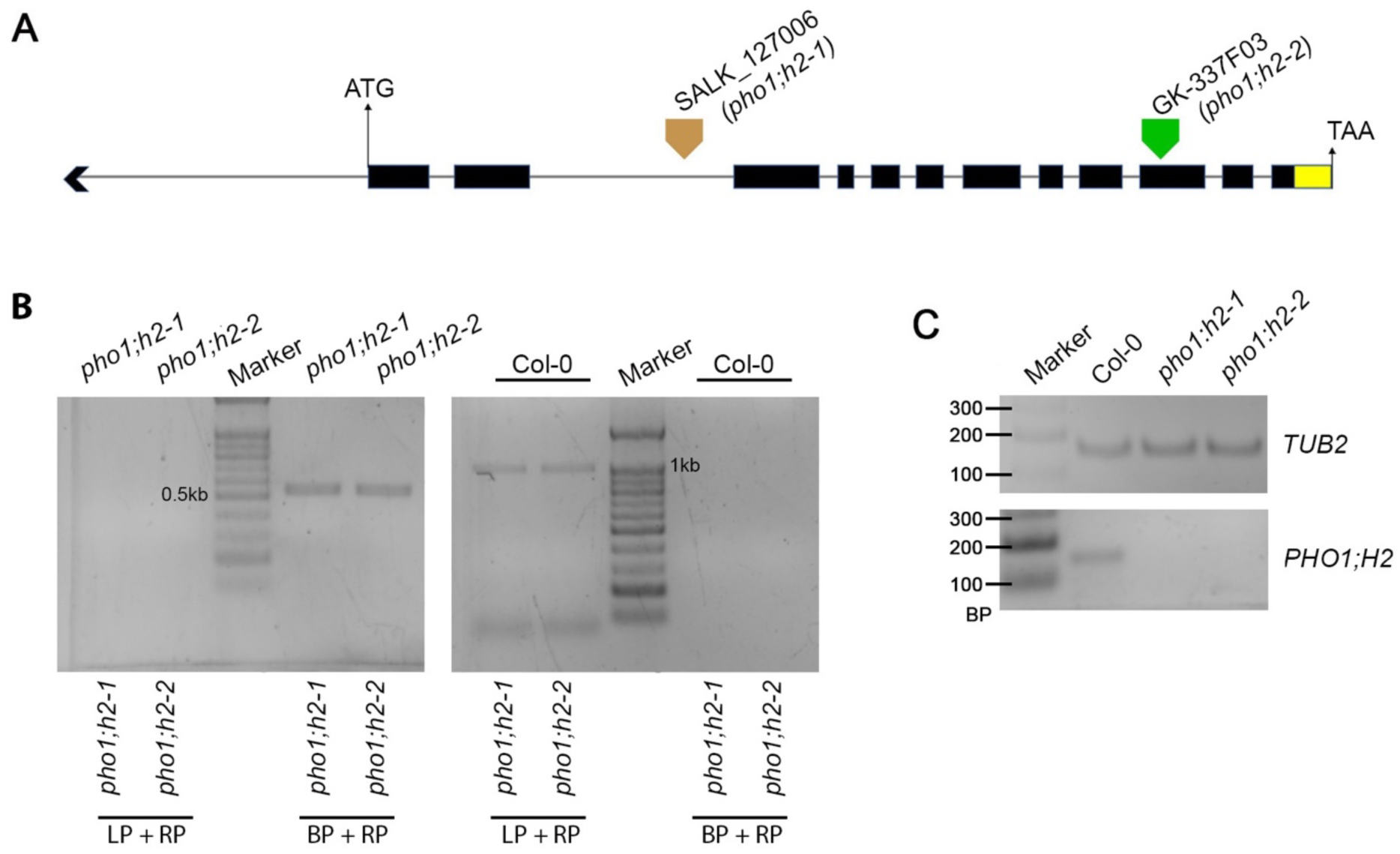
Identification of T-DNA insertion in mutants for *PHO1; H2* locus. (**A**) Identification of T-DNA insertion in mutants for PHO1;H2. Representation of PHO1;H2 locus. It has 12 exons (black bars) and 11 introns (lines). Arrows mark the T-DNA insertion positions. (**B**) PCR-based genotyping using gene-specific and T-DNA border-specific primers to confirm the T-DNA insertion mutant lines (*pho1;h2-1*, Salk_127006; *pho1;h2-2,* GK-337F03.11) in the *PHO1;H2* locus. (**C**) Molecular characterization of *pho1;h2* mutants by semiquantitative RT-PCR. Expression of *PHO1;H2* and *TUB2* in Col-0, *pho1;h2-1* and *pho1h2-2* genotypes, suggest that both *pho1;h2-1* and *pho1h2-2* are likely null alleles.

**Fig. S3.**
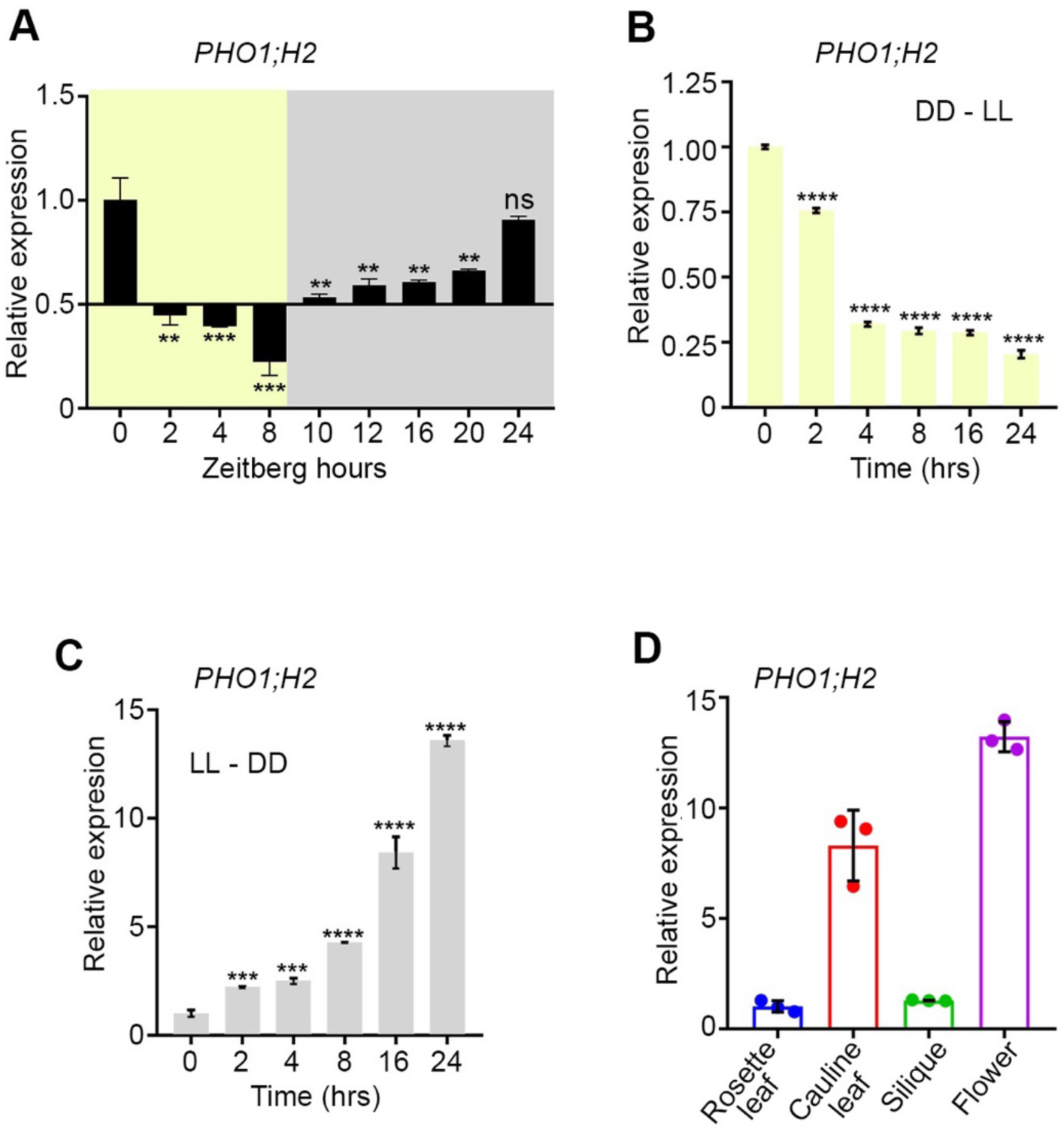
The PHO1;H2 gene expression is controlled by light and expressed in various tissues. **(A)** RT-qPCR analysis shows the diurnal regulation of *PHO1;H2* transcript accumulation in the wild-type. Tissue from six-day-old Col-0 seedlings grown at 22°C under SD was harvested at different ZT time points (0, 2, 4, 6, 8,10,16, 20, 24) spanning light and dark phase across the day. **(B-C)** RT-qPCR analysis to check the effect of light on *PHO1;H2* transcript accumulation. Seedling tissue was harvested at the respective time points and used for RT-qPCR analysis. Six-day-old continuous dark (DD) grown seedlings were shifted to WL for different time durations (0, 2, 4, 8,10,16, 24h) (B). Similarly, constant light (LL)-grown seedlings were shifted to darkness for different time durations (0, 2, 4, 8, 10, 16, 24 h) (C). The data represent mean±SD (n=3, biological replicates). Mean values were normalized to the transcript levels of *TUB2* internal control, and those values were further normalized to Col-0 (0 h time point) to get the fold-change. (**D**) Tissue-specific expression of *PHO1;H2* from Col-0 plants grown in WL at 22°C under long-day photoperiod at 22°C. Different aerial parts of Col-0, such as rosettes, cauline leaves, siliques, and flowers, were used for RT-qPCR analysis. The data represent the mean ± SD (n = 3, biological replicates). Mean values were normalized to the transcript levels of *TUB2* internal control, In A-D, the asterisk (*) represents the statistically significant differences from ZT0 or from the 0 h time point, as computed from the Student’s *t*-test (*p<0.05, **p<0.001, ***p<0.001 and ****p<0.0001).

**Fig. S4.**
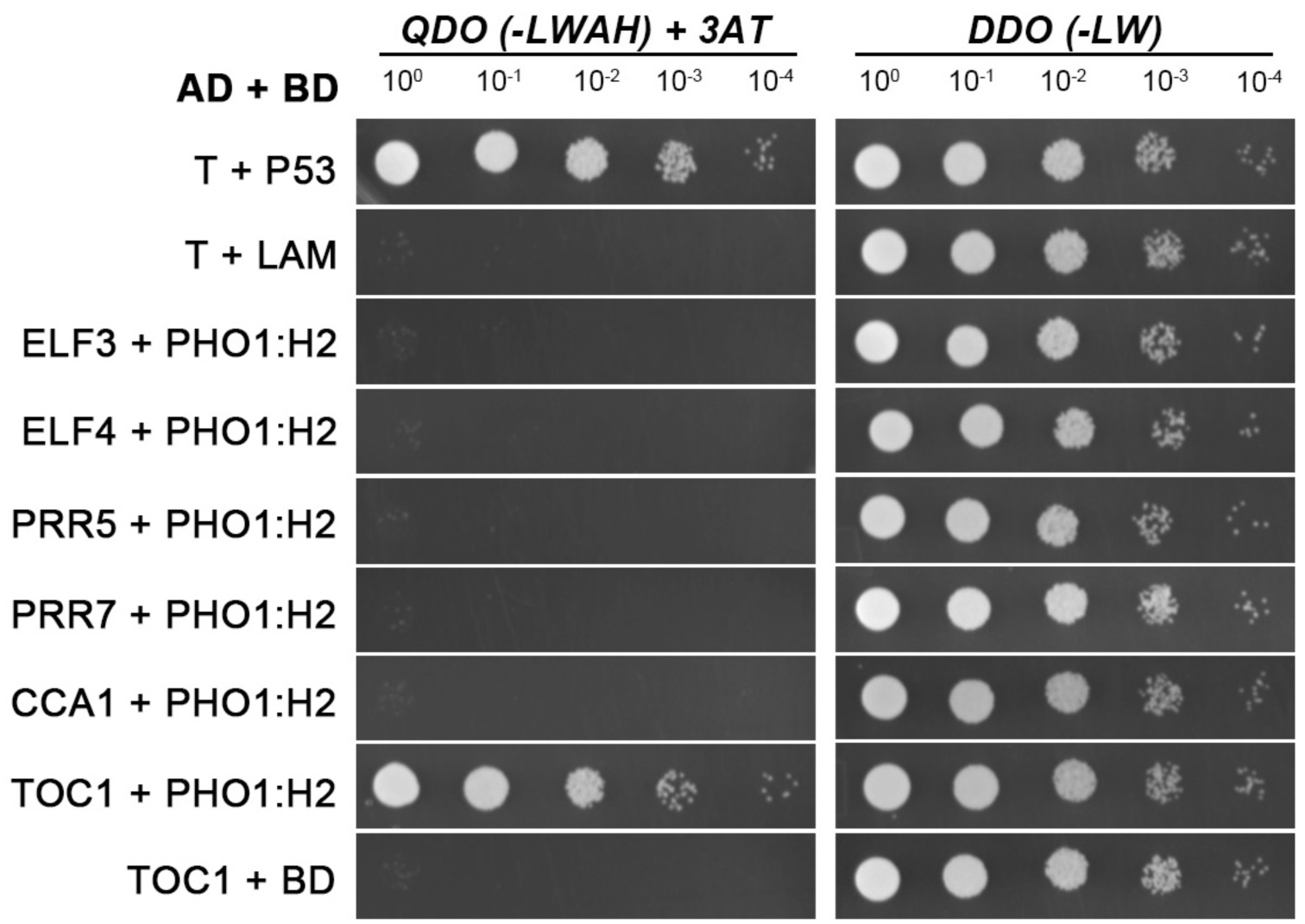
Yeast two-hybrid assay between PHO1;H2 and core circadian clock proteins. **(A)** Representative pictures of yeast two-hybrid interaction between *PHO1;H2* and circadian clock proteins ELF3, ELF4, PRR5, PRR7, CCA1, TOC1, and PIF4 with indicated dilution on double and quadruple dropout/ SD media with 3-AT.

**Fig. S5.**
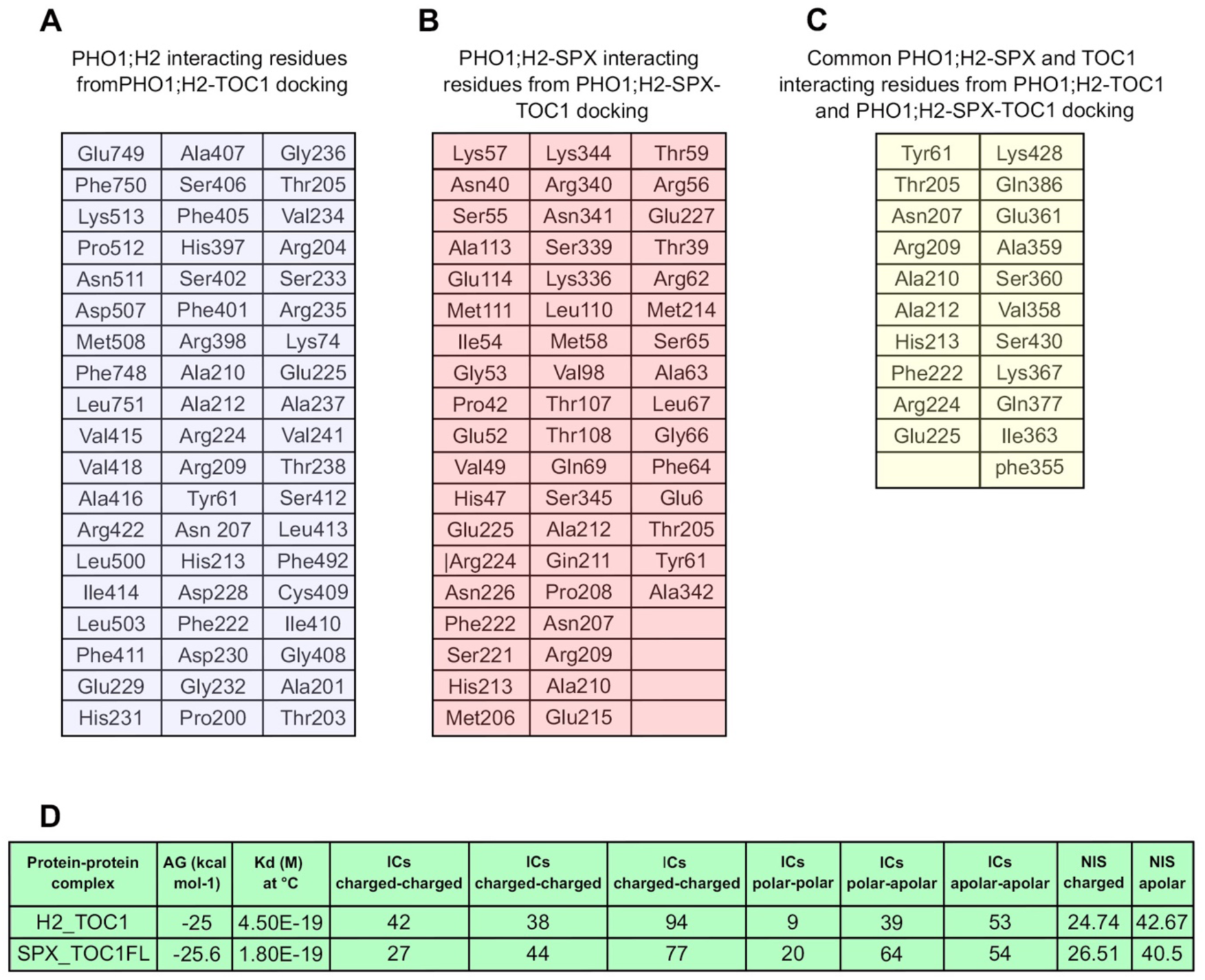
List of interacting amino acid residues from PHO1;H2-TOC1, and PHO1;H2-SPX-TOC1 docking. **(A)** Amino acid residues of PHO1;H2 predicted to participate in the interaction interface with TOC1, as identified by protein-protein docking analysis. The listed residues represent the key contact sites. **(B)** Predicted interacting residues within the SPX domain of PHO1;H2 derived from PHO1;H2-SPX and TOC1 docking analysis. These residues indicate that the SPX domain forms a major binding interface with TOC1 **(C)** Overlapping interacting residues common to both PHO1;H2-TOC1 and PHO1;H2-SPX-TOC1 docking complexes. The shared residues define a conserved interaction core, supporting a central role for the SPX domain in mediating PHO1;H2-TOC1 association **(D)** Comparative analysis of biding energetics and interfacial interaction profiles of FL length PHO1:H2 with TOC1 (H2_TOC1) and SPX with TOC1 (SPX_TOC1FL) complexes. Shown are Gibbs free energy(ΔG), predicted dissociation constant (K_d_), distribution of interfacial residues (IC_s_) by interaction type (Charged-Charged, Charged-polar, polar-polar, polar-apolar, and apolar-apolar) and non-interacting surface (NIS) contributions. The low (ΔG) and K_d_ values indicate strong and stable interactions.

**Fig. S6.**
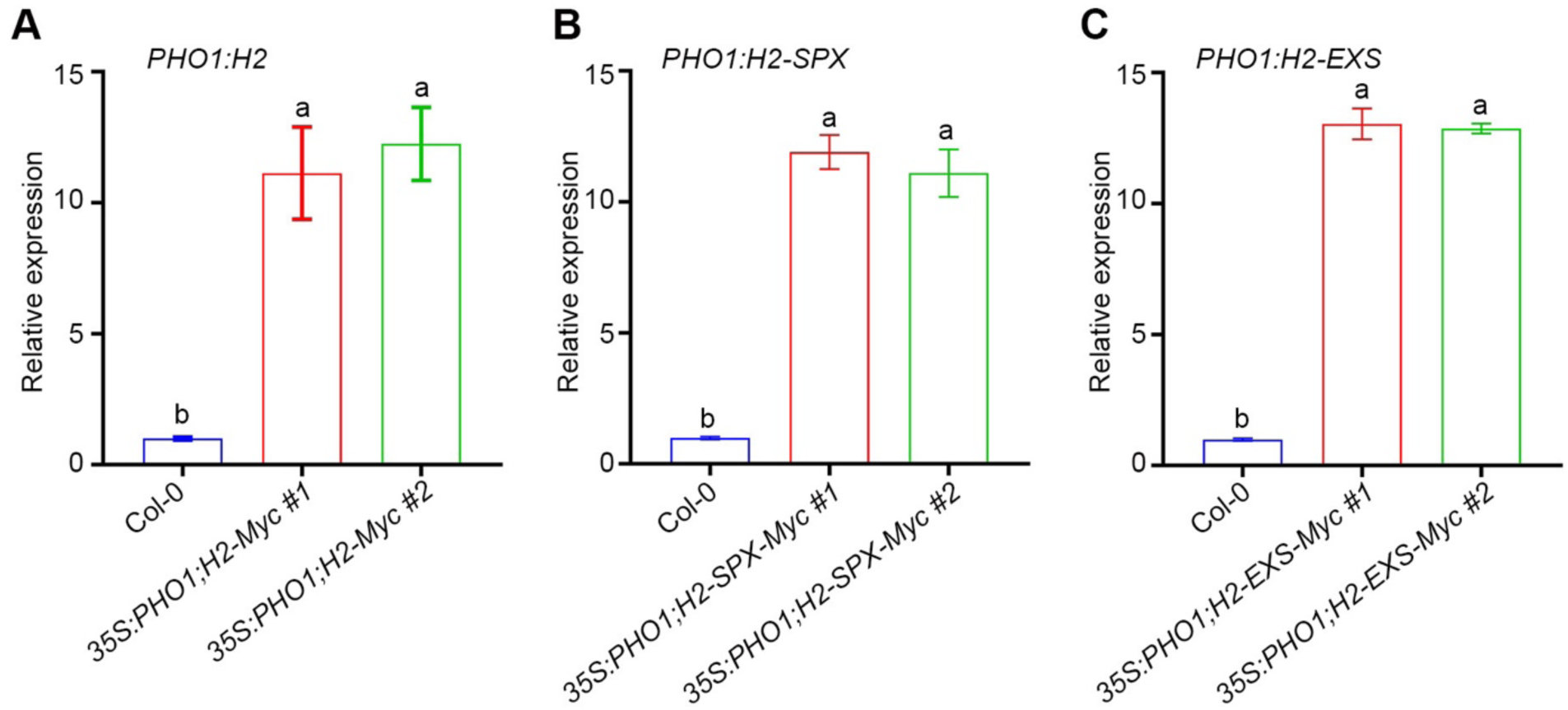
RT-qPCR analysis of transgenic lines overexpressing full-length, SPX and EXS domains of *PHO1;H2*. **(A-C)** Relative transcript levels of (A) *PHO1:H2*, (B) *PHO1:H2-SPX*, and (C) *PHO1:H2-EXS* in Col-0 background along with Col-0. Two independent *35S*-driven Myc-tagged overexpression lines for full-length (*35S:PHO1;H2-Myc #1, 35S:PHO1;H2-Myc #2*)*, SPX domain* (*35S:PHO1;H2-SPX-Myc #1*, *35S:PHO1;H2-SPX-Myc #2*), and EXS domain (*35S:PHO1;H2-EXS-Myc #1, 35S:PHO1;H2-EXS-Myc #1*) were used for checking the overexpression. Relative gene expression was calculated normalizing to Col-0. Bars represent mean±SD (n=3 biological replicates). *TUBULIN* was used as an internal normalizer. Different letters indicate statistically significant differences between genotypes as revealed by ANOVA followed by Tukey’s post-hoc HSD test (p<0.05).

**Fig. S7.**
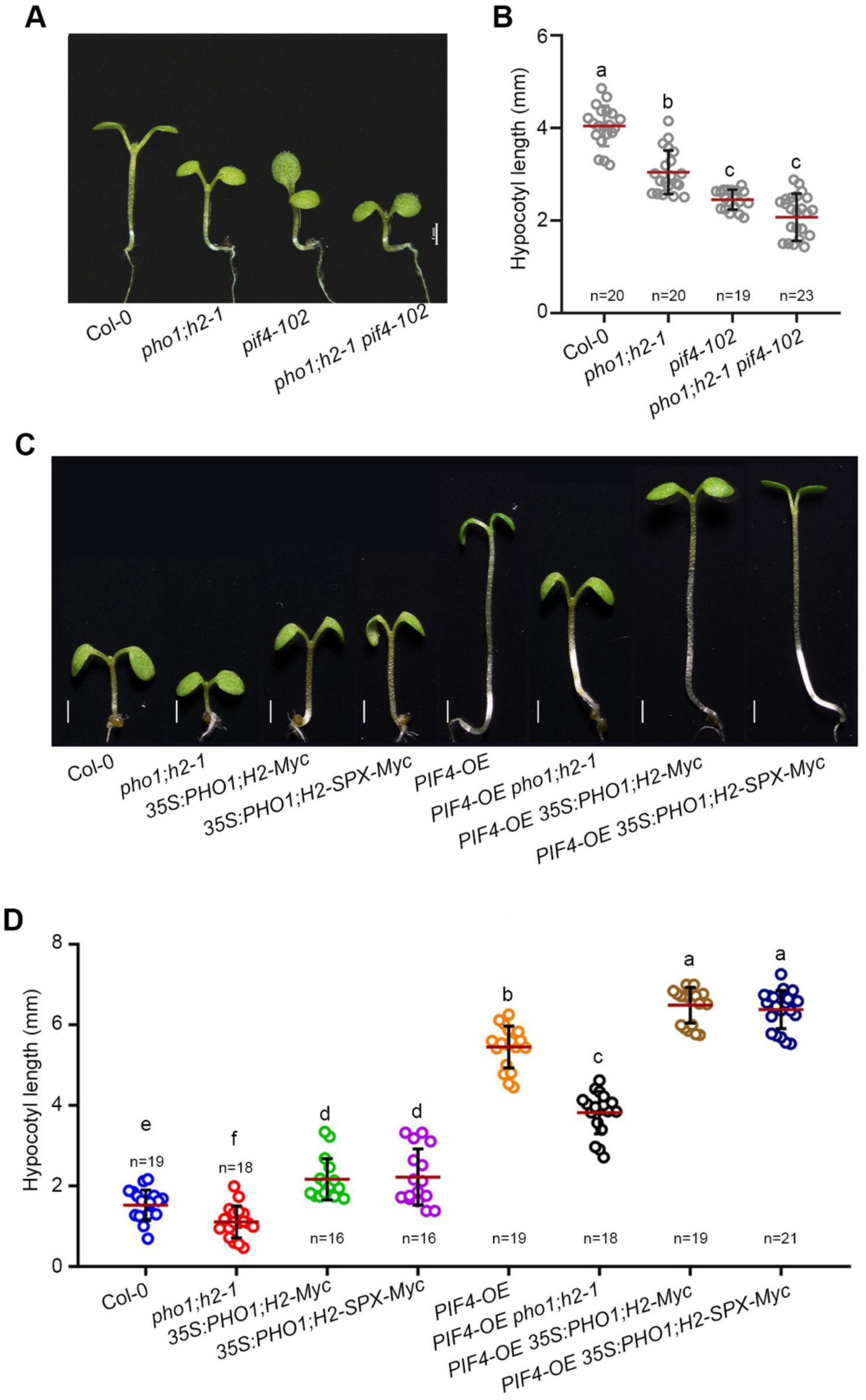
PHO1;H2 is essential for the optimal PIF4 function. **(A and B)** Representative hypocotyl images (A), and mean hypocotyl lengths (B) of six-day-old seedlings Col*-0, pho1;h2-1, pif4-101* and *pho1;h2-1 pif4-101* genotypes grown in WL (60 µmol m^-^²s^-^¹) at 22°C under SD. Scale, 1 mm. The data shown is the mean±SD (≥20) **(C)** Representative Hypocotyl images of 6-day-old seedlings Col-0, *pho1;h2-1, 35S:PHO1;H2-Myc, 35S:PHO1;H2-SPX-Myc , PIF4-OE, PIF4-OE pho1;h2-1, PIF4-OE 35S:PHO1;H2-Myc* and *PIF4-OE 35S:PHO1;H2-SPX-Myc* grown in WL (60 µmol m^-2^s^-1^) at 22°C under LD photoperiod. Scale, 1 mm. **(D)** Mean Hypocotyl measurements of Col-0, *pho1;h2-1, 35S:PHO1;H2-Myc, 35S:PHO1;H2-SPX-Myc , PIF4-OE, PIF4-OE pho1;h2-1, PIF4-OE 35S:PHO1;H2-Myc* and *PIF4-OE 35S:PHO1;H2-SPX-Myc* genotypes shown in A. The data shown is the mean±SD (n≥20). Small letters above the bars denote genotypes that significantly differ as revealed by one-way ANOVA followed by Tukey’s post-hoc HSD test (p<0.05). “*n*” indicates the number of seedlings measured.

**Fig. S8.**
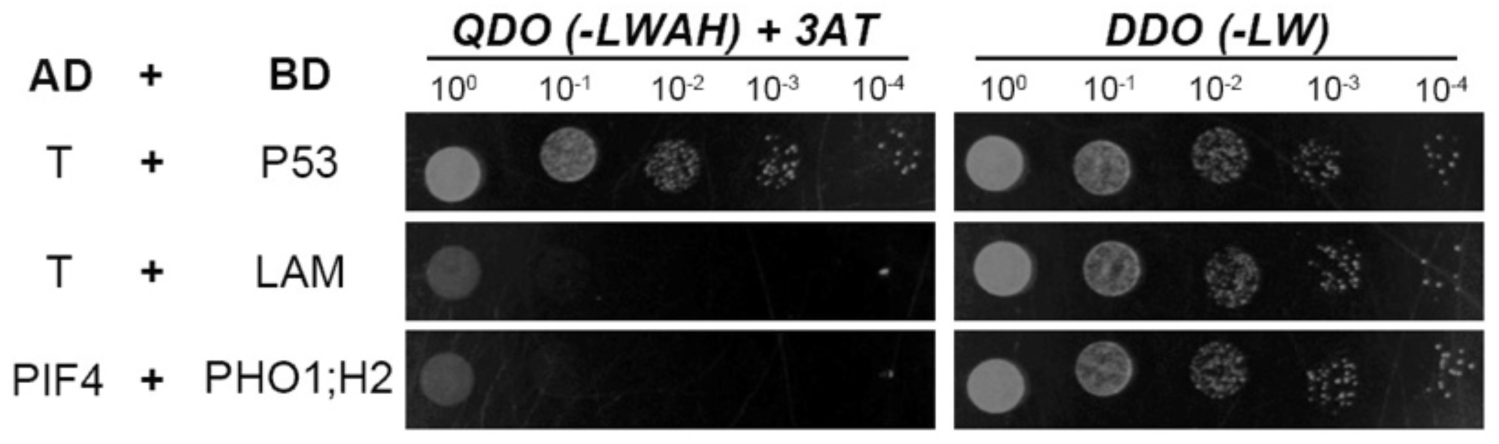
PHO1;H2 does not physically interact with PIF4. Representative pictures of yeast two-hybrid interaction between PHO1:H2 and PIF4 with indicated dilution on double [DDO] and quadruple [QDO] dropout/ SD media with 3-AT. AD-T and BD-P53 served as positive control, whereas AD-T with BD-LAM served as a negative control and failed to grow under selective conditions.

**Table S1.**
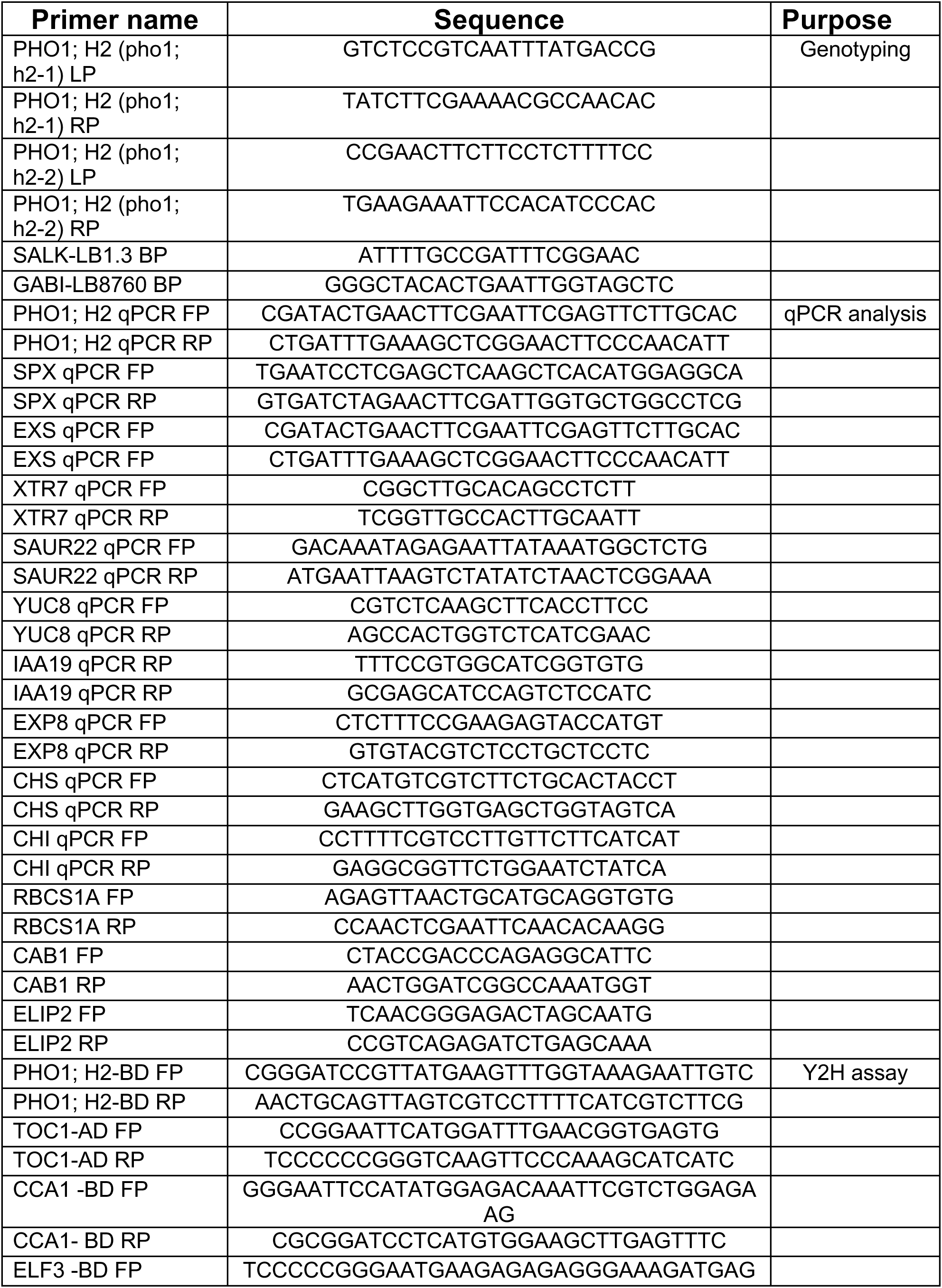
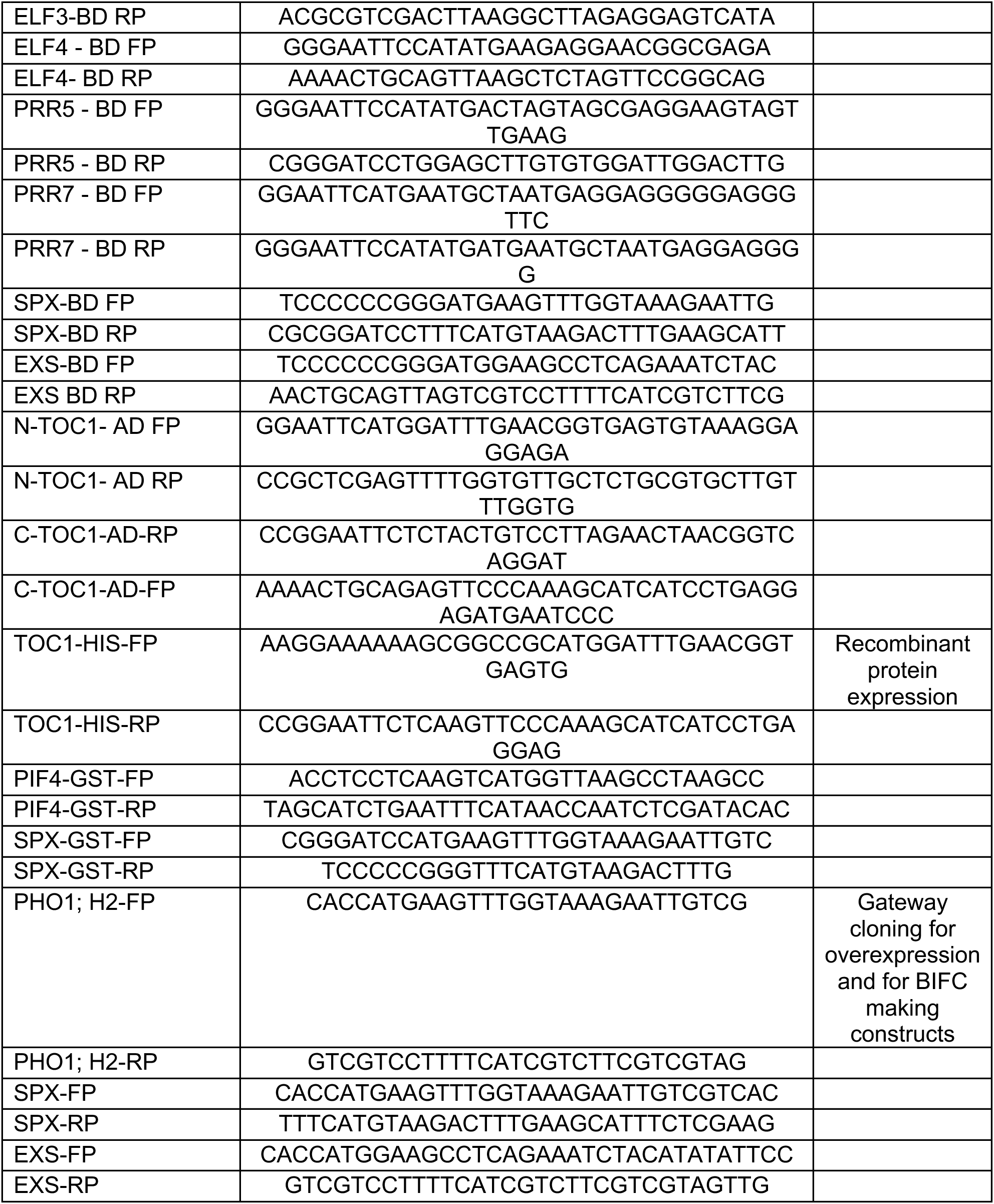
List of Oligonucleotides used in this study.

